# Large vs small genomes in *Passiflora*: the influence of the mobilome and the satellitome

**DOI:** 10.1101/2020.08.24.264986

**Authors:** Mariela Sader, Magdalena Vaio, Luiz Augusto Cauz-Santos, Marcelo Carnier Dornelas, Maria Lucia Carneiro Vieira, Natoniel Melo, Andrea Pedrosa-Harand

**Author notes:** Corresponding author, Universidade Federal de Pernambuco, Centro de Biociências, Departamento de Botânica, Laboratório de Citogenética e Evolução Vegetal, R. Prof. Moraes Rego, s/n, CDU. 50670-901 Recife PE Brazil, or 8352.

## Abstract

Repetitive sequences are ubiquitous and fast-evolving elements responsible for size variation and large-scale organization of plant genomes. Within *Passiflora* genus, a ten-fold variation in genome size, not attributed to polyploidy, is known. Here, we applied a combined *in silico* and cytological approach to study the organization and diversification of repetitive elements in three species of these genera representing its known range in genome size variation. Sequences were classified in terms of type and repetitiveness and the most abundant were mapped to chromosomes. We identified Long Terminal Repeat (LTR) retrotransposons as the most abundant elements in the three genomes, showing a considerable variation among species. Satellite DNAs (satDNAs) were less representative, but highly diverse between subgenera. Our results clearly confirm that the largest genome species (*Passiflora quadrangularis*) presents a higher accumulation of repetitive DNA sequences, specially Angela and Tekay elements, making up most of its genome. *Passiflora cincinnata*, with intermediate genome and from the same subgenus, showed similarity with *P. quadrangularis* regarding the families of repetitive DNA sequences, but in different proportions. On the other hand, *Passiflora organensis*, the smallest genome, from a different subgenus, presented greater diversity and the highest proportion of satDNA. Altogether, our data indicate that while large genome evolve by an accumulation of retrotransponsons, small genomes most evolved by diversification of different repeat types, particularly satDNAs.

**MAIN CONCLUSIONS:** While two lineages of retrotransposons were more abundant in larger *Passiflora* genomes, the satellitome was more diverse and abundant in the smallest genome.

## INTRODUCTION

Eukaryotic genomes are composed of a large amount of different classes of repetitive DNA sequences, either dispersed (mainly transposons and retrotransposons, as well as some protein-coding gene families) or arranged in *tandem* (ribosomal RNA, protein-coding gene families, and mostly satDNAs) (López-Flores & Garrido-Ramos, 2012; Biscotti et al. 2015). Transposable elements (TEs) represent up to 90% of the genome size, for example, 45% of the human genome (Lander et al. 2001), 52% of the opossum genome (Mikkelsen et al. 2007), or 85% of the maize genome (Schnable et al. 2009). Repetitive DNA sequences has been referred to as the repeatome (Goubert et al. 2015; Jouffroy et al. 2016; Pita et al. 2017; Hannan 2018). Repeat motifs can vary extensively in sequence and abundance (De Koning et al. 2011; Biscotti et al. 2015; Maumus & Quesneville 2016). Thus, transposable elements (retroelements and DNA transposons) and tandem repeats (satDNA and rDNA) have been postulated to have multiple roles in the genome, including genome stability, recombination, chromatin modulation and modification of gene expression (Biscotti et al. 2015). Apart from polyploidy, genome size variation in plants has been a consequence of increases and decreases in the number of these two types of sequences and may reflect different evolutionary strategies in speciation (Albach & Greilhuber 2004).

TEs are repeated DNA sequences, with the ability to move within genomes. Two classes of TEs are distinguished: Class I elements, or retrotransposons, use reverse transcriptase to copy an RNA intermediate into the host DNA. They are divided into Long Terminal Repeat (LTR) and non-LTR elements. Class II elements, or DNA transposons, use the genome DNA of the element itself as the template for transposition, either by a “cut and paste” mechanism, involving the excision and reinsertion of the DNA sequence of the element, or by using a rolling circle process or a virus-like process (Levin and Moran, 2011; Pritham, 2009). These two classes are subdivided into super-families and families based on their transposition mechanism, sequence similarities, structural features and phylogenetic relationships (Wicker et al. 2007; Neumann et al. 2019). The set of transposable elements in a genome is known as mobilome (Siefert 2009).

Satellite DNAs (satDNA) have been the most unknown part of genomes. Initially also considered as junk DNA, there is currently an increasing appreciation of its functional significance (Kidwell 2002; Garrido-Ramos 2017). SatDNA families accumulate mostly in the heterochromatin at different parts of the eukaryotic chromosomes, mainly in pericentromeric and subtelomeric regions, also spanning the functional centromere (Garrido-Ramos 2015). Their rapid evolution and constant homogenization (“concerted evolution”) (Hemleben et al. 2007) give rise to sequences that are genus-, species- or chromosome-specific (Rayburn and Gill 1986; Metzlaf et al. 1986; King 1995). This process generates divergence between species or reproductive groups (López-Flores and Garrido-Ramos 2012). The whole collection of satDNAs in a genome is also known as satellitome (Ruiz-Ruano et al. 2016).

The study of repetitive DNA and its impact in genome size evolution has significantly progressed since the introduction of next generation sequencing (NGS) technologies (Margulies et al. 2005) associated with new and improved bioinformatic analyses. For example, the clustering procedure of genome sequence reads based on similarity has been improved by employing graph-based methods (Novák et al. 2010). This approach has been efficient in the identification and characterization of repeat elements in several organisms (Aversano et al. 2015; Derks et al. 2015; Wolf et al. 2015, Ribeiro et al 2019, Van Lume et al 2019, Gaiero et al 2019; McCann et al 2020).

The genus *Passiflora* L. belongs to the family Passifloraceae Juss. ex Kunth, which is a member of the Malpighiales order (Judd et al. 2015). *Passiflora* is a large and morphologically variable genus and it includes 575 species distributed in the tropical and subtropical regions of America, Africa and Asia (Ulmer and MacDougal, 2004). The species of *Passiflora* show a substantial variation in chromosome size and number, with different basic chromosome numbers (*x* = 6, 9, and 12) for different subgenera or clades (Melo and Guerra 2003; Hansen et al. 2006; Sader et al. 2019a). Variation in genome sizes also has been reported for the genus (Yotoko et al. 2011). Considering all data available for genome size of 62 species, comprising around 10% of the genus, the difference between the largest and smallest genomes is currently as high as 10 times [0.212 pg in *P. organensis* Gardner*, (Decaloba* subgenus); and 2.68 pg in *P. quadrangularis* L., (*Passiflora* subgenus)] (Yotoko et al. 2011; Souza et al. 2004). A recent diversification in the subgenus *Passiflora* (Miocene) was associated to chromosome number change from *n* = 6 to *n* = 9 and an increase in genome size. Polyploidy was restricted to few lineages and was not associated with species diversification or genome size variation. Thus, dysploidy together with genome size increase could have acted as the main drivers in the evolution of *Passiflora* (Sader et al. 2019a).

Two recent works describe the repetitive fraction of the genome of the “yellow passion-fruit”, *P. edulis* Sims. (Costa et al. 2019; Pamponét et al. 2019). A total of 250 different TE sequences were identified (96% Class I, and 4% Class II), corresponding to ~19% of the *P. edulis* draft genome, assembled *de novo* from Illumina NGS reads (Araya et al. 2017). TEs were found preferentially in intergenic spaces (70.4%), but also overlapping with genes (30.6%). As in most plant species, the highest proportion of the genome is represented by LTR retrotransposons totalling 53% of the genome (Pamponét et al. 2019). Ribosomal DNA (5S and 35S) accounted for 1% of the genome, and the lowest proportion was also observed for satDNAs, reaching less than 0.1% (Costa et al. 2019). A phylogenetic inference of the reverse transcriptase domain of the LTR-retrotransposons and insertion time analysis showed that the majority (95.9%) of the LTR-retrotransposons were recently inserted into the *P. edulis* genome (< 2.0 Mya). With the exception of the Athila lineage, all LTR-retrotransposons were transcriptionally active. In addition, some lineages appeared to be conserved in wild *Passifora* species (Costa et al. 2019). In the light of this information, the aim of this work was to understand the cause of the large variation of genome size in the genus *Passiflora*. For this, three species were chosen, the smallest genome species, the largest genome species and a medium-sized genome. Their genomes were sequenced with low coverage and reads were clustered by sequence similarities to recognize and characterize the most abundant sequences at chromosomal level.

## MATERIAL AND METHODS

### Plant material

The materials used in this work included *P. organensis* Gardner (2*n* = 2*x* = 12), *P. cincinnata* Maxwell (2*n* = 2*x* = 18) (access CPI54) and *P. quadrangularis* L. (2*n* = 2*x* = 18) plants (access QPE68) from “Banco Ativo de Germoplasma, Embrapa Semiárido”. Although *P. organensis* is considered synonymous of *P. porophylla* Vell. (The Plant List, http://www.theplantlist.org/), the individuals from Paranapiacaba (São Paulo, Brazil) correspond to *P. organensis* sensu stricto (Cauz-Santos, sinbiota 22603). The plants used for sequencing and cytogenetics were maintained in the Experimental Garden of the Federal University of Pernambuco, Recife, Brazil.

### Next generation sequencing, processing data, and clustering analysis

DNA isolation from leaves of *P. cincinnata* and *P. quadrangularis* plants was carried out according to Weising (2005). Sequencing was performed on an Illumina HiSeq using 250bp paired-end reads in BGI Group (Hong Kong, China). Sequencing of *P. organensis* was performed on an Illumina HiSeq (250bp) in the Earlham Institute (Norwich, UK). All sequences were filtered using a cut-of value of 20 and a 90% of bases equal or above this value.

The similarity-based clustering analysis was performed using the RepeatExplorer pipeline at the Elixir-Cerit server (https://repeatexplorer-elixir.cerit-sc.cz) (Novák et al. 2010, 2013). We performed two different analyses: 1) an individual analysis for each species, with a larger coverage (Table 1) to better characterize all TE families using automatic option sampling in RepeatExplorer; 2) a comparative analysis, using reads from the libraries of the three species. For the latter, interlaced reads for each species were identified with a prefix and then concatenated. We used 561,853 total reads. Combined, the repeats identified for each species represented 41,542 reads for *P. organensis* (0.05×), 235,288 reads for *P. cincinnata* (0.04×), and 285,022 for *P. quadrangularis* (0.03×) of the total genome for each species (Table 1). Coverage was calculated as follow: coverage = (r × l)/g, whereas r corresponds to number of reads used in our analysis, l to read length and g to haploid genome size in bp. The concatenated dataset was run in the same pipeline but using the comparative analysis option.

**Table 1.** Genome sizes and sequencing read number and coverage for the analysed *Passiflora* species

|  | *P. organensis* | *P. cincinnata* | *P. quadrangularis* |
| --- | --- | --- | --- |
| $2n =$ | 12 | 18 | 18 |
| Genome size (1C) | 0.212 pg | 1.420 pg | 2.680 pg |
|  | 207.34 Mbp | 1,388.76 Mbp | 2,621.04 Mbp |
| Individual clustering reads | 600,000 | 494,343 | 1,044,658 |
| Coverage | 0.72× | 0.09× | 0.1× |
| Comparative clustering reads | 41,542 | 235,288 | 285,022 |
| Coverage | 0.05× | 0.04× | 0.03× |

Resulted clusters with genomic percentage above 0.01% were further manually examined to characterize the most abundant repetitive families. Unclassified clusters were analysed by similarity searches using BLASTN and BLASTX against non-redundant protein sequences public databases (https://blast.ncbi.nlm.nih.gov/Blast.cgi). The TAREAN tool from Repeat Explorer was applied for satellite DNA repeat identification (Novák et al.2017). Further examination was performed for each cluster based on graph layout, those with a ringlike shape were selected and analysed as potential satDNA. Repeat composition was calculated excluding clusters of organelle DNA probably representing extranuclear DNA from chloroplast and mitochondria.

### Satellite DNA characterization

In order to perform a high-throughput analysis of satellite DNA and detect as many families as possible in the genome, we used the satMiner pipeline, a toolkit for mining and analysing satDNA families (https://github.com/fjruizruano/satminer). For this we followed the protocol suggested by Ruiz-Ruano et al. (2016). Briefly, the protocol consists of reads quality trimming with Trimmomatic and then clustering a selection of 2□×□200,000 reads with RepeatExplorer. We identified the satellite repeats based on the typical tandem repeat graph-layout (i.e., spherical or ring-shaped), and confirmed their repetitive structures and monomer length with the dotplot tool in Geneious R8.1 software (Kearse et al. 2012). Subsequently, we used DeconSeq software to filter out reads showing homology with all previously identified clusters. Then, using a sample of the remaining reads, we started an additional round performing a new RepeatExplorer clustering duplicating the number of reads for each new run. The protocol was repeated for several runs until no new satellite repeats were identified or no more reads were available.

For final characterization and annotation, we looked for homology among found satDNAs. We performed two different analyses: 1) we used RepeatMasker v4.05 to aligned each satellite sequence to the rest. 2) To know if all satDNA found in *P. organensis* were present in the other two species, we used ‘map to reference’ tool in Geneious Prime 2019.0.4 to search for these sequences against all reads (12,222,646 reads in *P. quadrangularis*; 6,758,428 reads in *P. cincinnata* and, 112,367,646 reads in *P. organensis*). To investigate the degree of homology among each of the characterized satDNAs, we considered that monomeric sequences with 50 - 80% similarity belonged to different families of the same superfamily of satDNAs. Also, sequences with 80 - 95% similarity were variants of the same family (ie. subfamily), and those showing > 95% similarity were considered to be variants of the same monomer, as proposed by Ruiz-Ruano et al. (2016). We also employed their nomenclature rules for satDNA: the name begin with species abbreviation (three letters) followed by the term “Sat”, then a catalog number in order of decreasing abundance and finally the consensus monomer length.

We determined the abundance for each variant by using the ‘map to reference’ tool in Geneious Prime 2019.0.4 and we calculated the relative abundance by dividing the number of mapped reads by the total number of reads. In order to amplify the annotated satDNAs by PCR, we aligned satDNA monomers to get a consensus sequence and selected the most conserved region to design primers with Primer3 tool (Untergasser 2012) in Geneious R8.1 software (Kearse et al. 2012). Primers were designed facing towards ensuring to minimize the distance between them or even overlapping them up to three bp at the 5’ end, when necessary.

In addition, to check if one satDNA originated from the IGS region of the rDNA, we assembled the complete 35S rDNA sequence of the three species with NOVOPlasty (Dierckxsens et al. 2016) using a random selection of 10 million read pairs. As a seed reference, we used the 5.8S rDNA sequence from *Passiflora edulis* species obtained from GenBank (accession number MF327245).

### Phylogeny of Gypsy-Tekay

With the purpose of better understand the dynamics of Ty3/gypsy-Tekay retrotransposons in the three species, we analysed these elements in more detail using similarity searches against all Gypsy elements identified in the retrotransposon protein domain database – ReXdb (Neumann et al. 2019). The Gypsy-Tekay Integrase protein domains were extracted from full-length REs using the RepeatExplorer platform (Novak et al. 2010). Further, nucleotide sequences were translated in all possible reading frames and the resulting peptides were aligned together with a specific set of integrases from Ty3/gypsy elements using MAFFT (Katoh & Standley, 2013) with “Auto” option. The alignment was used to construct phylogenetic trees using Neighbor Joining Tree Protein using Geneious Prime 2019.0.4 (http://www.geneious.com, Kearse et al. 2012).

### Molecular cytogenetic techniques

Root tips obtained from plants growing in pots were pretreated with 2 mM 8-hydroxyquinoline for 4,5 h at 10°C, fixed in ethanol–acetic acid (3:1 v/v), and stored in fixative at −20°C. Root tips were digested using a solution containing 2% cellulase and 20% pectinase (w/v) for 90 min at 37 °C and chromosome preparations were performed according to Carvalho and Saraiva (1993). For each species, the repetitive DNA for the probes were isolated by PCR. The PCR mix contained template DNA (25 ng), 1× PCR buffer (20 mM Tris-HCl pH 8.4, 50 mM KCl), 0.25 mM of each dNTP, 0.5 μM of each primer and homemade Taq DNA polymerase in a total volume of 50 μl. The PCR program consisted of 35 cycles, each with 1 min denaturation at 94°C, 1 min annealing 55–57°C (depending on the primer pair used), 1 min extension at 72°C. PCR products were purified by precipitation and one microgram each was labeled by *nick translation* (Invitrogen or Roche Diagnostics) with Cy3-dUTP (GE) or Alexa 448-5-dUTP (Life Technologies). The 25-28S, 5.8S, and 18S rDNA clone (p*Ta*71) from *Triticum aestivum* (Gerlach and Bedbrook 1979), labeled with digoxigenin-11-dUTP (Roche) was used to localize 35S rDNA sites.

The fluorescent *in situ* hybridization (FISH) procedure applied to mitotic chromosomes was essentially the same as previously described (Fonsêca et al. 2010). Hybridization mix consisted of 50% (v/v) formamide, 10% (w/v) dextran sulfate, 2× SSC, and 2-5 ng/μl probe. The slides were denatured for 5 min at 75°C and hybridized for 24 h at 37°C. The final stringency was 76%. Images were captured in an epifluorescence Leica DMLB microscope using the Leica QFISH software and Leica DMLB microscope using Leica Las-AF software. For final processing, images were artificially coloured using the Adobe Photoshop software version 10.0 and uniformly adjusted for brightness and contrast only.

## RESULTS

### Genomic Composition

The repeat fraction of each individual genome was characterized using around 0.1× coverage for *P. cincinnata* and *P. quadrangularis* and 0.72× for the small *P. organensis* genome (Tables 1 and 2; Fig1.).

**Table 2.**
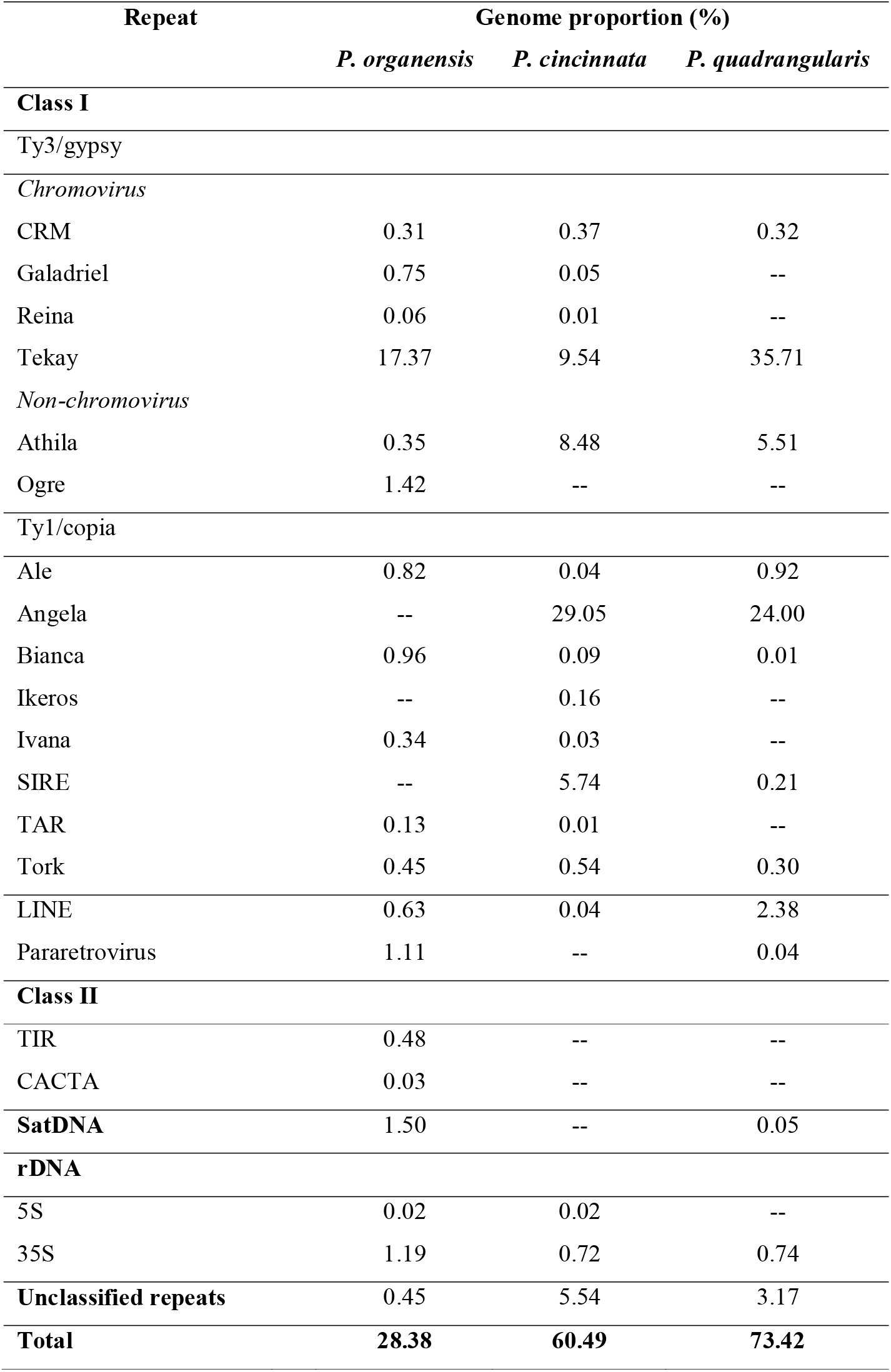
Genome proportion (%) of repetitive sequences identified in the individual RepeatExplorer analyses of *Passiflora organensis* (*Decaloba* subgenus), and *P. cincinnata* and *P. quadrangularis* (*Passiflora* subgenus).

| Repeat | Genome proportion (%) |  |  |
| --- | --- | --- | --- |
|  | *P. organensis* | *P. cincinnata* | *P. quadrangularis* |
| **Class I** |  |  |  |
| Ty3/gypsy |  |  |  |
| *Chromovirus* |  |  |  |
| CRM | 0.31 | 0.37 | 0.32 |
| Galadriel | 0.75 | 0.05 | -- |
| Reina | 0.06 | 0.01 | -- |
| Tekay | 17.37 | 9.54 | 35.71 |
| *Non-chromovirus* |  |  |  |
| Athila | 0.35 | 8.48 | 5.51 |
| Ogre | 1.42 | -- | -- |
| Ty1/copia |  |  |  |
| Ale | 0.82 | 0.04 | 0.92 |
| Angela | -- | 29.05 | 24.00 |
| Bianca | 0.96 | 0.09 | 0.01 |
| Ikeros | -- | 0.16 | -- |
| Ivana | 0.34 | 0.03 | -- |
| SIRE | -- | 5.74 | 0.21 |
| TAR | 0.13 | 0.01 | -- |
| Tork | 0.45 | 0.54 | 0.30 |
| LINE | 0.63 | 0.04 | 2.38 |
| Pararetrovirus | 1.11 | -- | 0.04 |
| **Class II** |  |  |  |
| TIR | 0.48 | -- | -- |
| CACTA | 0.03 | -- | -- |
| **SatDNA** | 1.50 | -- | 0.05 |
| **rDNA** |  |  |  |
| 5S | 0.02 | 0.02 | -- |
| 35S | 1.19 | 0.72 | 0.74 |
| **Unclassified repeats** | 0.45 | 5.54 | 3.17 |
| **Total** | **28.38** | **60.49** | **73.42** |

**Fig. 1.**
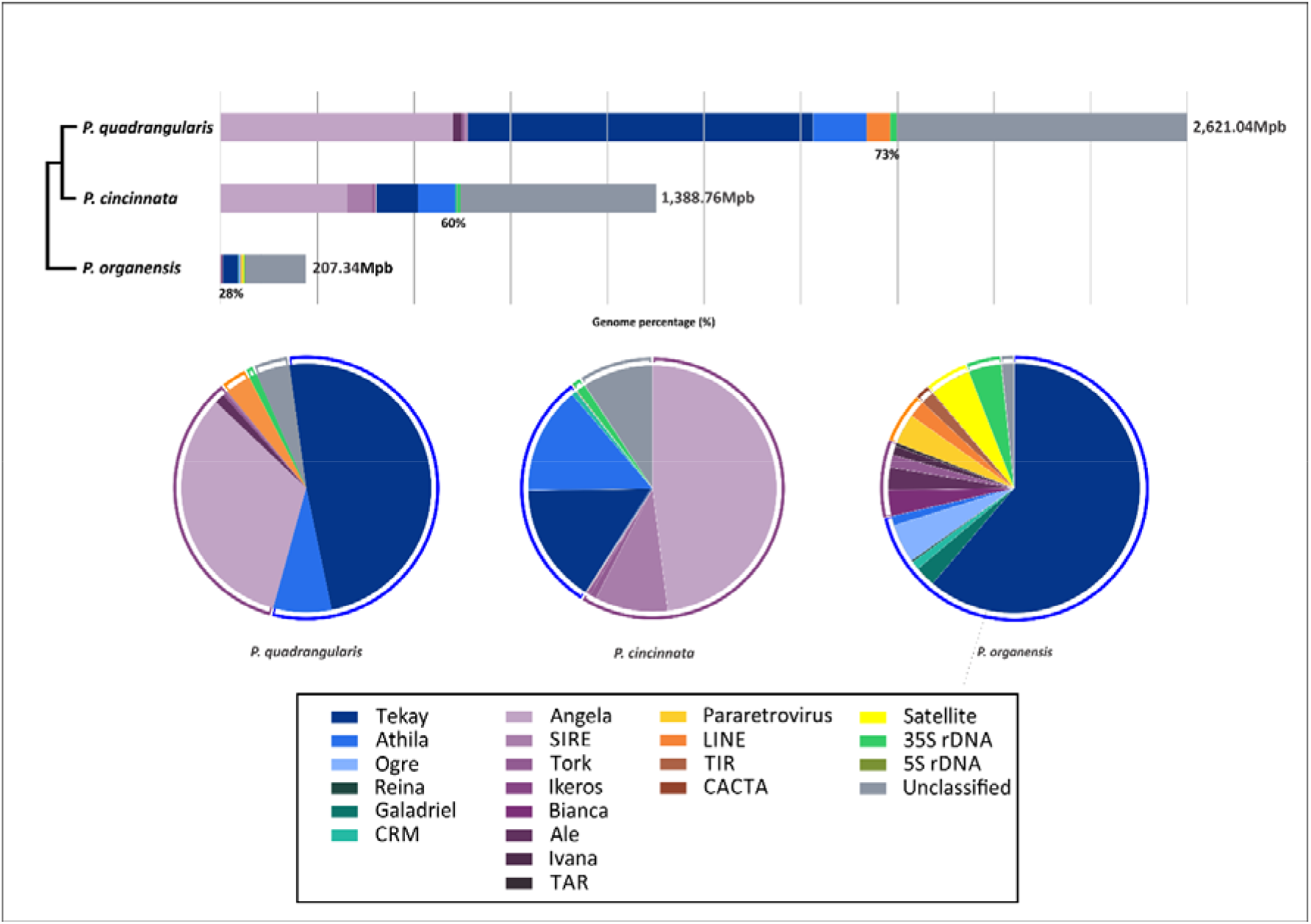
Summary of the composition of the repetitive fraction originated from individual clustering analysis in the three *Passiflora* genomes and their evolutionary relationship based on (Sader et al. 2019a).

#### P. organensis

The 299 clusters that corresponded to at least 0.01% of the genome (from 84 up to 11,269 reads) contained 722,865 reads corresponding to 28.38% of the genome. The superfamily Ty3/gypsy from the LTR retrotransposonswas the most abundant, dominated by Tekay lineage (17.37%) and Galadriel (0.75), against 2.70% of Ty1/copia, mainly Bianca. Satellite DNA corresponded to 1.50% of the genome. One cluster corresponded to 5S rDNA (0.02%) and three clusters to 35S rDNA (1.9%) (Table 2).

#### P. cincinnata

All 252 clusters (from 49 up to 10,774 reads) contained 455,846 reads corresponding to 60.49% of the genome. The Ty1/copia superfamily was the most abundant, corresponded to 35.69%, (mainly Angela lineage), followed by Ty3/gypsy retrotransposons (18.45%, from which 9.54% of Chromovirus Tekay, while 8.48% Gypsy/non-Chromovirus Athila). LINEs, 35S rDNA and 5S rDNA represent 0.04%, 0.72% and 0.025% of the genome respectively, while satDNAs were not identified among the 252 most abundant clusters (Table 2).

#### P. quadrangularis

All 161 clusters (from 106 up to 34,517 reads) contained 994,022 reads corresponding to 73.42% of the genome. Ty3/gypsy was the most abundant superfamily, corresponding to 41.54%, with Tekay (35.71%) and Athila (5.51%) as the main lineages. Ty1/copia elements represented 25.44%, with Angela (24%), Ale (0.92%), Tork (0.30%), SIRE (0.21%) and Bianca (0.01%) contributing unevenly. LINEs represented 2.38%, while the 35S rDNA corresponded to 0.74% of the genome (Table 2).

### Comparative analyses of DNA repetitive sequences

Comparative repetitive DNA analysis resulted in 267 clusters. The most abundant and shared element among the three species was Ty3/gypsy-Tekay. Although this element is present in the three species, most of the clusters are shared by species of the *Passiflora* subgenus only. There are also species-specific clusters, such as cluster 26, present in *P. quadrangularis* and cluster 29, which is the only one found in *P. organensis* (Fig. 2). Ty1/copia-Angela was very abundant in species of the *Passiflora* subgenus (*P. cincinnata* and *P. quadrangularis*) but absent in *P. organensis*. Finally, Ty1/copia-SIRE clusters were also shared by both species of the subgenus *Passiflora*, although in greater abundance in *P. cincinnata* (Fig. 2).

**Fig. 2.**
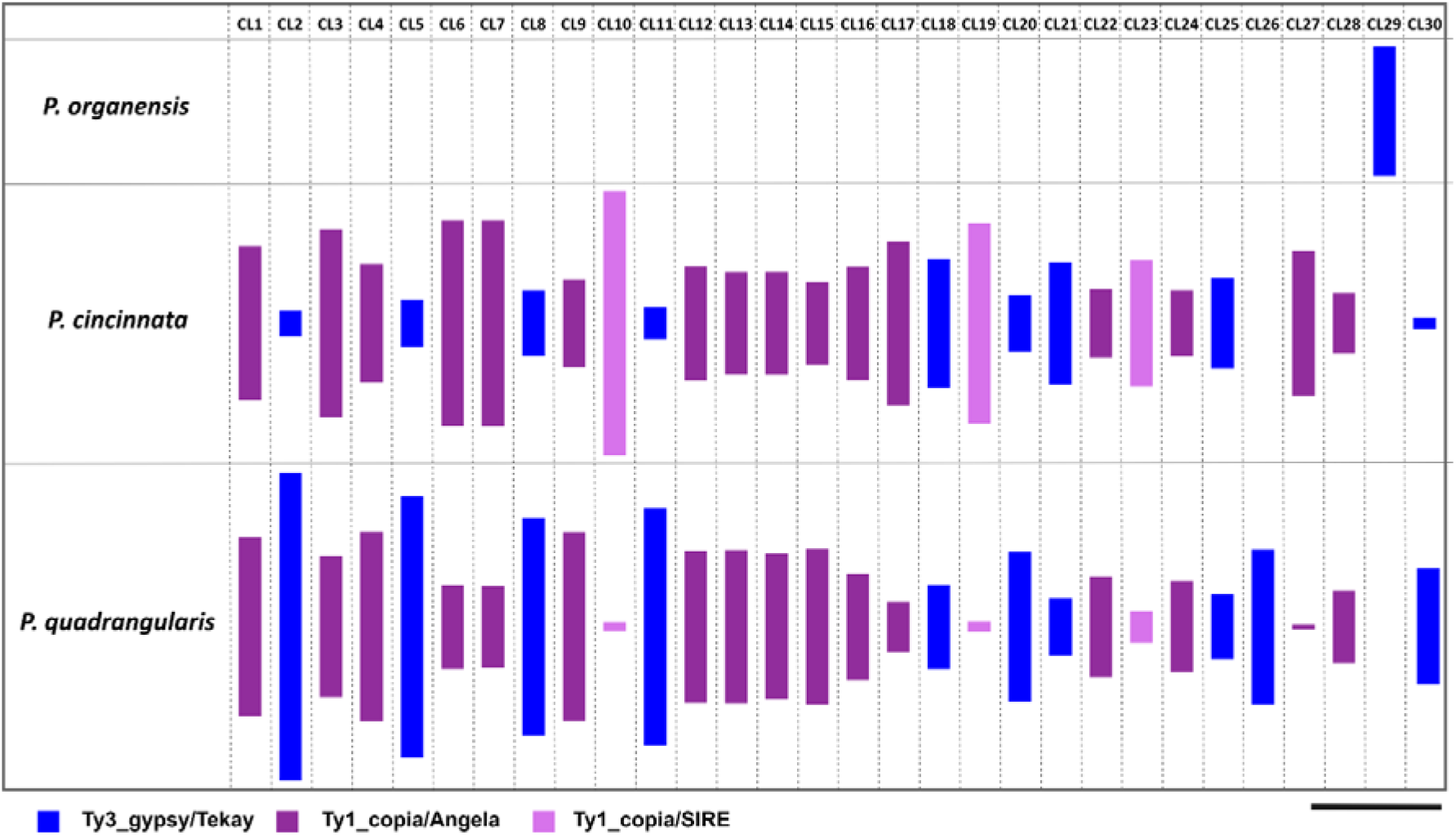
Representation of genomic abundances of the 30 largest clusters originated from comparative clustering analysis in *Passiflora*. The height of rectangle is proportional to the number of reads in a given cluster. Bar = 4,000 reads.

### Phylogeny of Ty3/gypsy-Tekay

Because the Ty3/gypsy-Tekay lineage was shared and abundant in the three sampled species, it was further analysed using similarity searches against the Gypsy elements from the retrotransposon protein domain database – ReXdb (Neumann et al. 2019) to better understand its divergence. Most of the Tekay elements recovered were incomplete or truncated. The phylogenetic reconstruction of the integrase domain of full-length Tekay elements revealed that similar elements are shared among species (Fig. 3). The first and second clades were shared by the three species. The third clade is shared by the *Passiflora* subgenera species (*P. cincinnata* and *P. quadrangularis*) and the fourth lineage has grouped only *P. organensis* elements.

**Fig. 3.**
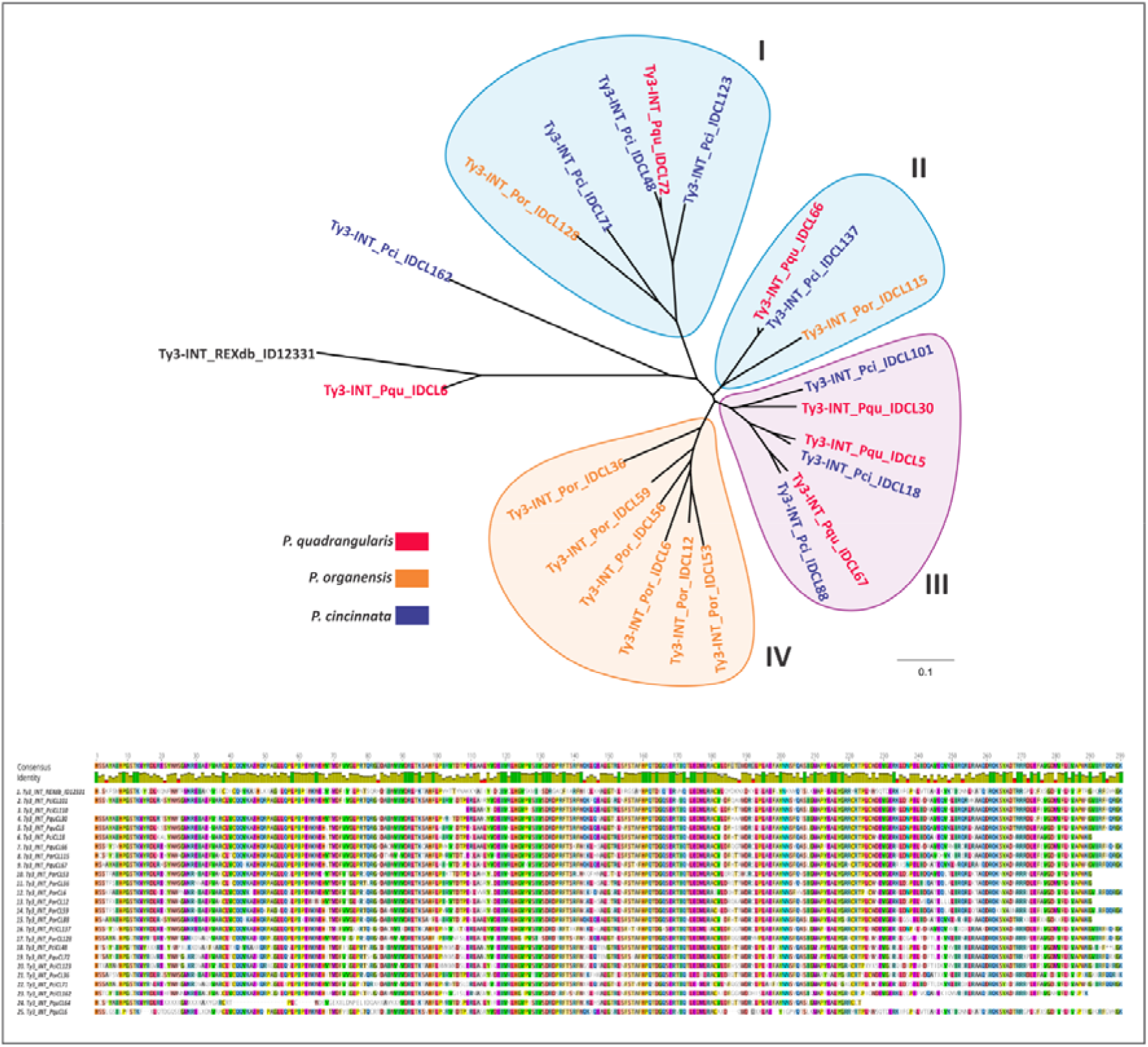
Phylogenetic relationship of Ty3/gypsy-Tekay elements of three *Passiflora* species based on the integrase domain (INT) aminoacids. Sequences used for comparison were retrieved from RexDB (Neumann et al. 2019). Details of integrase domain (INT) species alignment from MAFFT.

### Satellitome in Passiflora

Because no satDNA was identified among the most abundant repeats in the *Passiflora* subgenus species, we used the toolkit satMiner to find repeats that were in lower abundance in the genome of both *Passiflora* subgenus species but also in *P. organensis*. We performed nine iterations in *P. organensis* and only two in *P. cincinnata* and *P. quadrangularis* because no reads remained after these runs for both species. In total, we found 46 different satDNA families: 38 (16 in the first two runs) for *P. organensis*, 6 for *P. quadrangularis* (with only 1 in the second run), and 2 for *P. cincinnata* (none in the second run). Repeat unit lengths range between 52 and 3,998 bp (Table 3). The A+T content of the consensus satDNA sequences varied between 38.2% and 76.2% among families, with 60.9% median value, indicating a slight bias towards A+T rich satDNA.

**Table 3.**
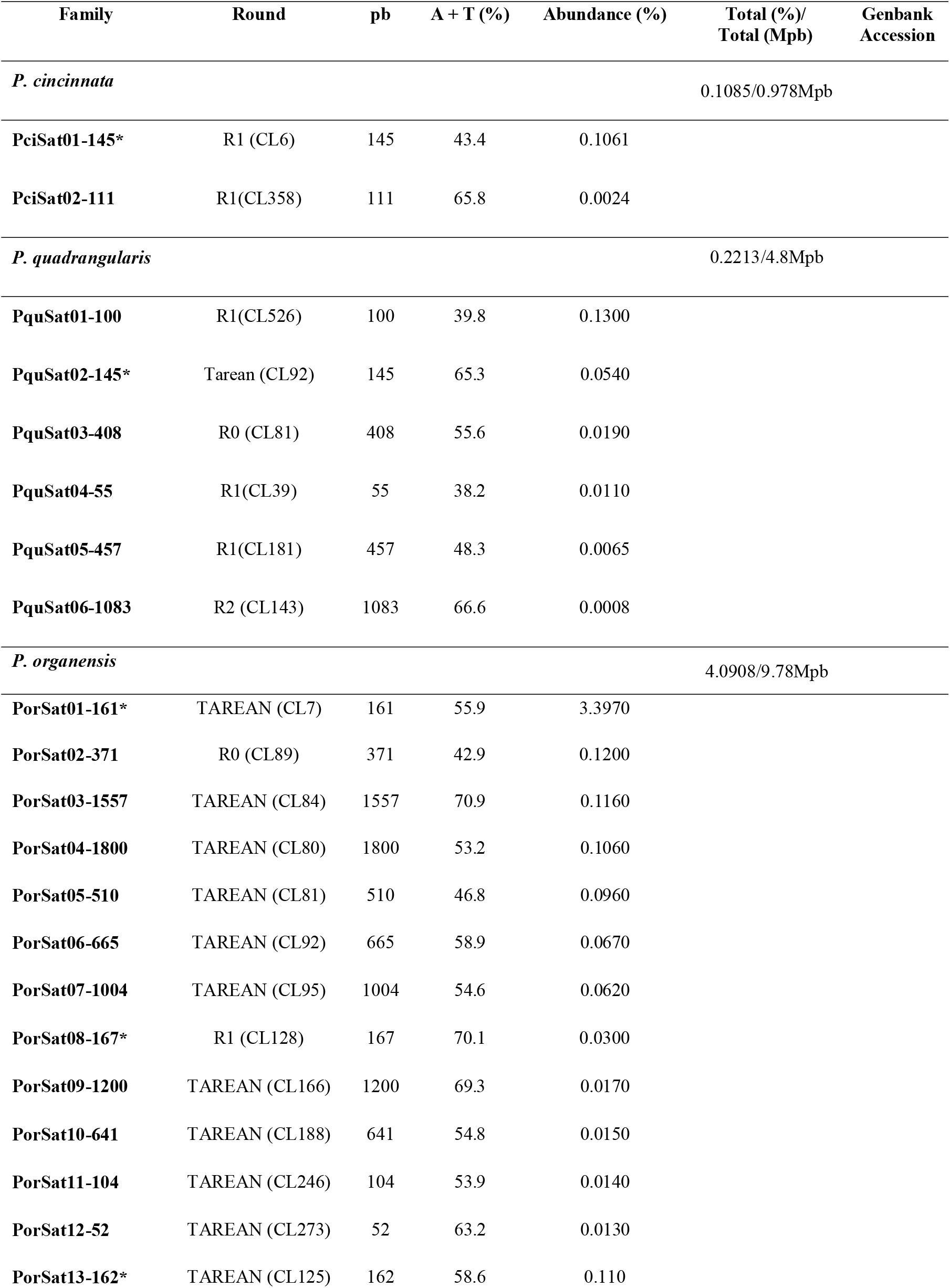
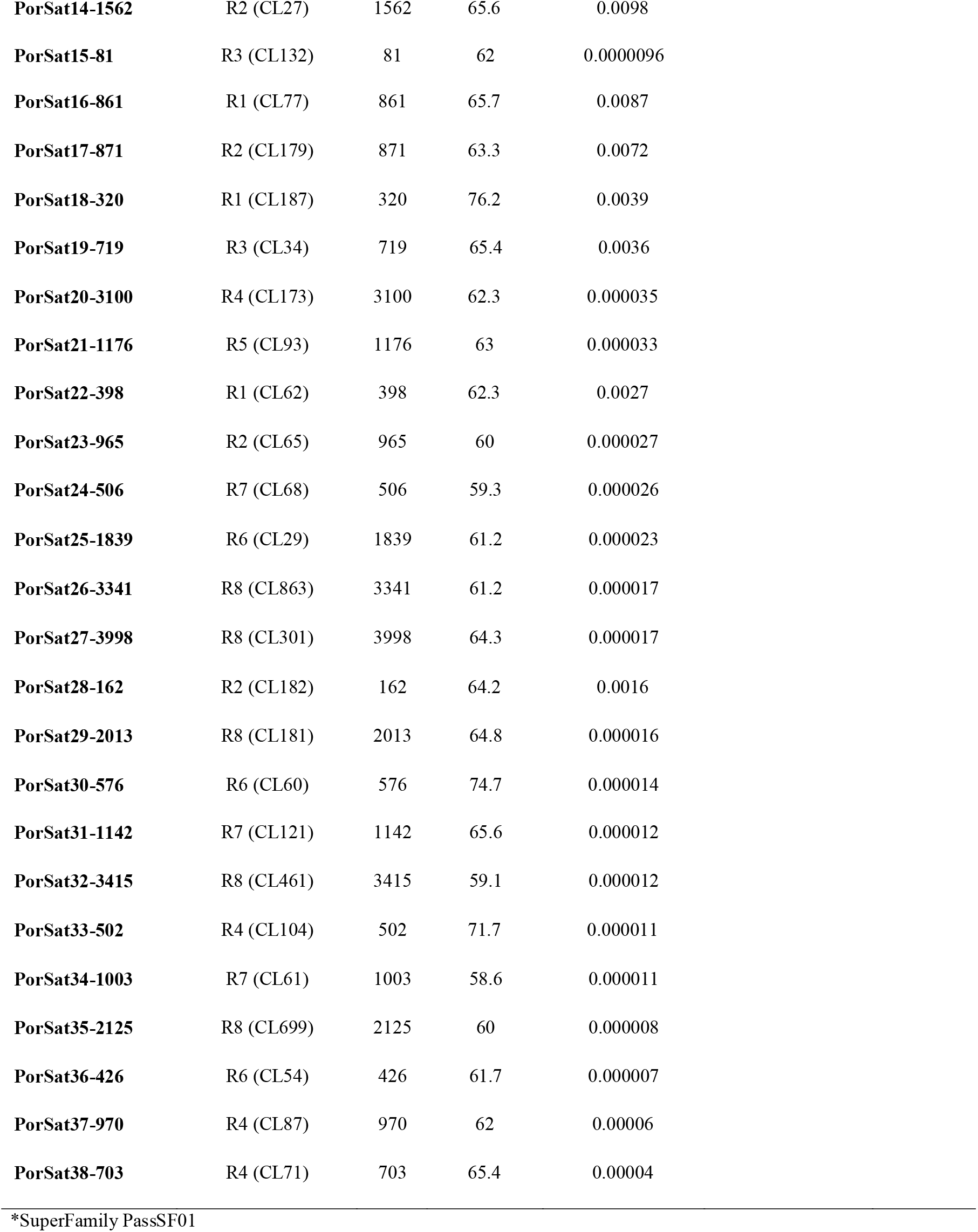
SatDNAs identified using the RepeatExplorer and satMiner pipelines for *P. cincinnata* and *P. quadrangularis* (*Passiflora* subgenus), and *Passiflora organensis (Decaloba* subgenus), showing length (nt), A + T content (%), abundance (%), and Genbank accession number.

| Family | Round | pb | A + T (%) | Abundance (%) | Total (%)/<br>Total (Mpb) | Genbank<br>Accession |
| --- | --- | --- | --- | --- | --- | --- |
| *P. cincinnata* |  |  |  |  | 0.1085/0.978Mpb |  |
| **PciSat01-145\*** | R1 (CL6) | 145 | 43.4 | 0.1061 |  |  |
| **PciSat02-111** | R1(CL358) | 111 | 65.8 | 0.0024 |  |  |
| *P. quadrangularis* |  |  |  |  | 0.2213/4.8Mpb |  |
| **PquSat01-100** | R1(CL526) | 100 | 39.8 | 0.1300 |  |  |
| **PquSat02-145\*** | Tarean (CL92) | 145 | 65.3 | 0.0540 |  |  |
| **PquSat03-408** | R0 (CL81) | 408 | 55.6 | 0.0190 |  |  |
| **PquSat04-55** | R1(CL39) | 55 | 38.2 | 0.0110 |  |  |
| **PquSat05-457** | R1(CL181) | 457 | 48.3 | 0.0065 |  |  |
| **PquSat06-1083** | R2 (CL143) | 1083 | 66.6 | 0.0008 |  |  |
| *P. organensis* |  |  |  |  | 4.0908/9.78Mpb |  |
| **PorSat01-161\*** | TAREAN (CL7) | 161 | 55.9 | 3.3970 |  |  |
| **PorSat02-371** | R0 (CL89) | 371 | 42.9 | 0.1200 |  |  |
| **PorSat03-1557** | TAREAN (CL84) | 1557 | 70.9 | 0.1160 |  |  |
| **PorSat04-1800** | TAREAN (CL80) | 1800 | 53.2 | 0.1060 |  |  |
| **PorSat05-510** | TAREAN (CL81) | 510 | 46.8 | 0.0960 |  |  |
| **PorSat06-665** | TAREAN (CL92) | 665 | 58.9 | 0.0670 |  |  |
| **PorSat07-1004** | TAREAN (CL95) | 1004 | 54.6 | 0.0620 |  |  |
| **PorSat08-167\*** | R1 (CL128) | 167 | 70.1 | 0.0300 |  |  |
| **PorSat09-1200** | TAREAN (CL166) | 1200 | 69.3 | 0.0170 |  |  |
| **PorSat10-641** | TAREAN (CL188) | 641 | 54.8 | 0.0150 |  |  |
| **PorSat11-104** | TAREAN (CL246) | 104 | 53.9 | 0.0140 |  |  |
| **PorSat12-52** | TAREAN (CL273) | 52 | 63.2 | 0.0130 |  |  |
| **PorSat13-162\*** | TAREAN (CL125) | 162 | 58.6 | 0.110 |  |  |

|  |  |  |  |  |
| --- | --- | --- | --- | --- |
| **PorSat14-1562** | R2 (CL27) | 1562 | 65.6 | 0.0098 |
| **PorSat15-81** | R3 (CL132) | 81 | 62 | 0.0000096 |
| **PorSat16-861** | R1 (CL77) | 861 | 65.7 | 0.0087 |
| **PorSat17-871** | R2 (CL179) | 871 | 63.3 | 0.0072 |
| **PorSat18-320** | R1 (CL187) | 320 | 76.2 | 0.0039 |
| **PorSat19-719** | R3 (CL34) | 719 | 65.4 | 0.0036 |
| **PorSat20-3100** | R4 (CL173) | 3100 | 62.3 | 0.000035 |
| **PorSat21-1176** | R5 (CL93) | 1176 | 63 | 0.000033 |
| **PorSat22-398** | R1 (CL62) | 398 | 62.3 | 0.0027 |
| **PorSat23-965** | R2 (CL65) | 965 | 60 | 0.000027 |
| **PorSat24-506** | R7 (CL68) | 506 | 59.3 | 0.000026 |
| **PorSat25-1839** | R6 (CL29) | 1839 | 61.2 | 0.000023 |
| **PorSat26-3341** | R8 (CL863) | 3341 | 61.2 | 0.000017 |
| **PorSat27-3998** | R8 (CL301) | 3998 | 64.3 | 0.000017 |
| **PorSat28-162** | R2 (CL182) | 162 | 64.2 | 0.0016 |
| **PorSat29-2013** | R8 (CL181) | 2013 | 64.8 | 0.000016 |
| **PorSat30-576** | R6 (CL60) | 576 | 74.7 | 0.000014 |
| **PorSat31-1142** | R7 (CL121) | 1142 | 65.6 | 0.000012 |
| **PorSat32-3415** | R8 (CL461) | 3415 | 59.1 | 0.000012 |
| **PorSat33-502** | R4 (CL104) | 502 | 71.7 | 0.000011 |
| **PorSat34-1003** | R7 (CL61) | 1003 | 58.6 | 0.000011 |
| **PorSat35-2125** | R8 (CL699) | 2125 | 60 | 0.000008 |
| **PorSat36-426** | R6 (CL54) | 426 | 61.7 | 0.000007 |
| **PorSat37-970** | R4 (CL87) | 970 | 62 | 0.00006 |
| **PorSat38-703** | R4 (CL71) | 703 | 65.4 | 0.00004 |

\*SuperFamily PassSF01

For final characterization, we looked for homology among all satDNAs, using RepeatMasker to align each satellite sequence to the rest. Sequence comparison between repeat units of the 46 satDNAs monomers detected homology between PorSat01-161 and PorSat13-162 (84%) and PorSat08-167 (56.3%), representing different families. PciSat01-145 and PquSat02-145 shared 89.3% of sequences homology, and can be considered subfamilies of the same family. These five satDNA showed 54.5% of homology and were grouped into superfamily 1 (PassSF01) (Online Resource 1; Table 3). In the second approach, we search if all satDNA found in *P. organensis* were present in the other two species. This analysis showed that almost all *P. organensis* satellites are exclusive, except PorSat20-3100 that was also found in *P. cincinnata* (0.0000001%) and *P. quadrangularis* (0.000001%), while PorSat37-970 was presented in *P. quadrangularis* (0.04%), but not detected in *P. cincinnata* genome.

### *Chromosomal Localization of most abundant repeats in* Passiflora

Aiming to determine the chromosomal location of the most abundant TE families, Ty1/copia-Angela and Ty3/gypsy-Tekay, we prepared probes containing the integrase domain from *P. quadrangularis* for FISH. We analyze Ty3/gypsy-Tekay on large genomes only. Angela and Tekay elements, abundant in the subgenus *Passiflora*, were dispersed in *P. quadrangularis* and *P. cincinnata* chromosomes, with a brighter signal at proximal regions in some chromosomes (Fig. 4).

**Fig. 4.**
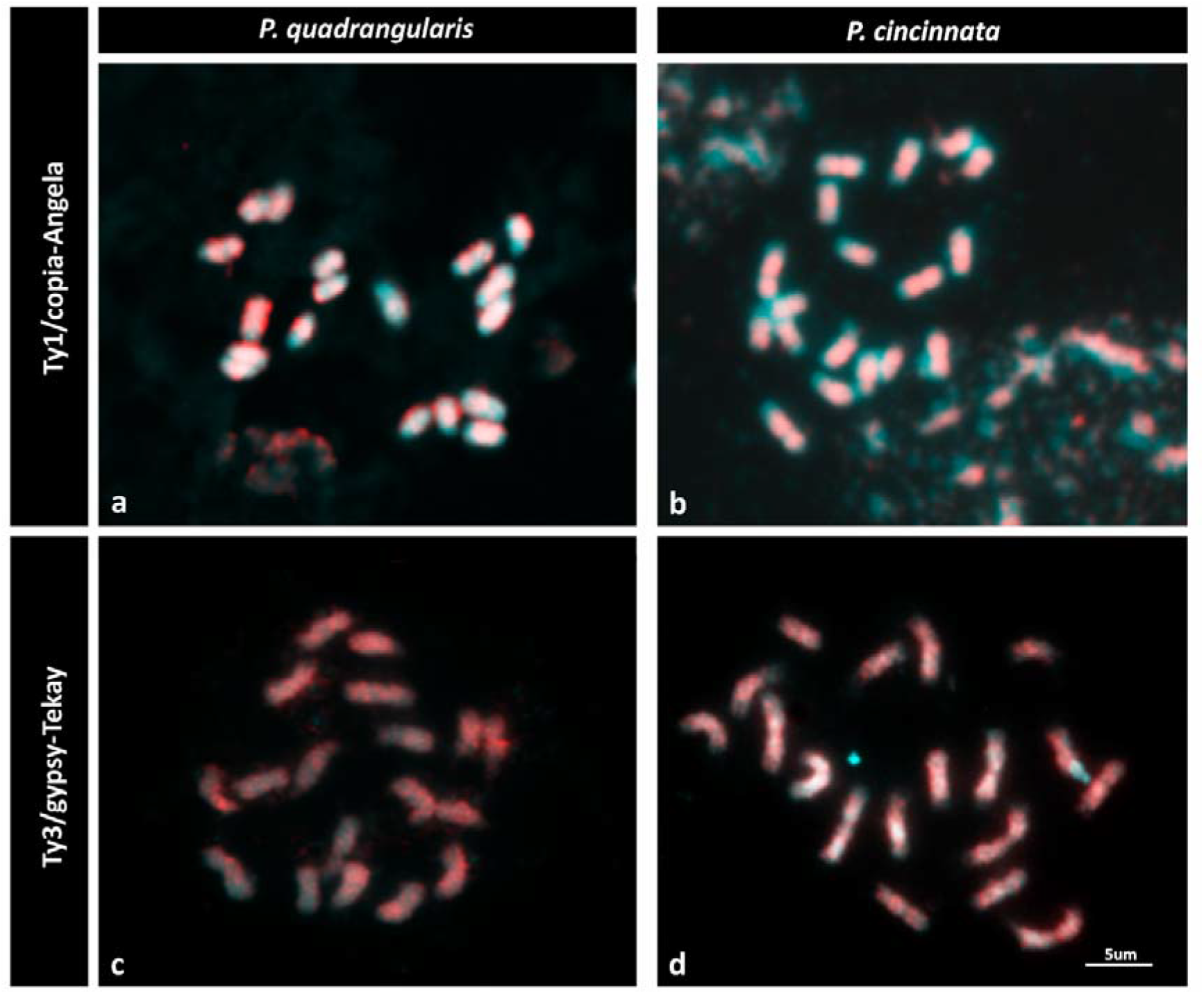
Chromosomal distribution of the Ty1/copia-Angela and Ty3/Gypsy-Tekay element in *Passiflora* genomes. (**a, c**) *Passiflora quadrangularis*, (**b, d**) *P. cincinnata*.

The most abundant satDNA families localized to the chromosomes of the three species. We used satellites recovered by TAREAN and in the first rounds of satMiner (R0 and R1) as probe. The DNAsat family PquSat02-145 (PassSF01) hybridized predominantly in subterminal sites, but also dispersedly, in most chromosomes of *P. quadrangularis* and *P. cincinnata* (Figs 5a and c). Satellite PquSat01-100 showed six terminal signals, as well as dispersed signals in chromosomes of *P. quadrangularis* (Fig. 5b), and four terminal signals in *P. cincinnata* (Fig. 5d), although no satDNA with homology to PquSat01-100 was identified in *P. cincinnata*. The strong signals co-localized with the 35S rDNA sites (Fig. 5d). The other satDNAs from *P. quadrangularis* and *P. cincinnata* (PquSat03-408, PquSat04-55, PquSat05-457, PquSat06-1083 and PciSat02-111) showed variable dispersed patterns (data not shown).

**Fig. 5.**
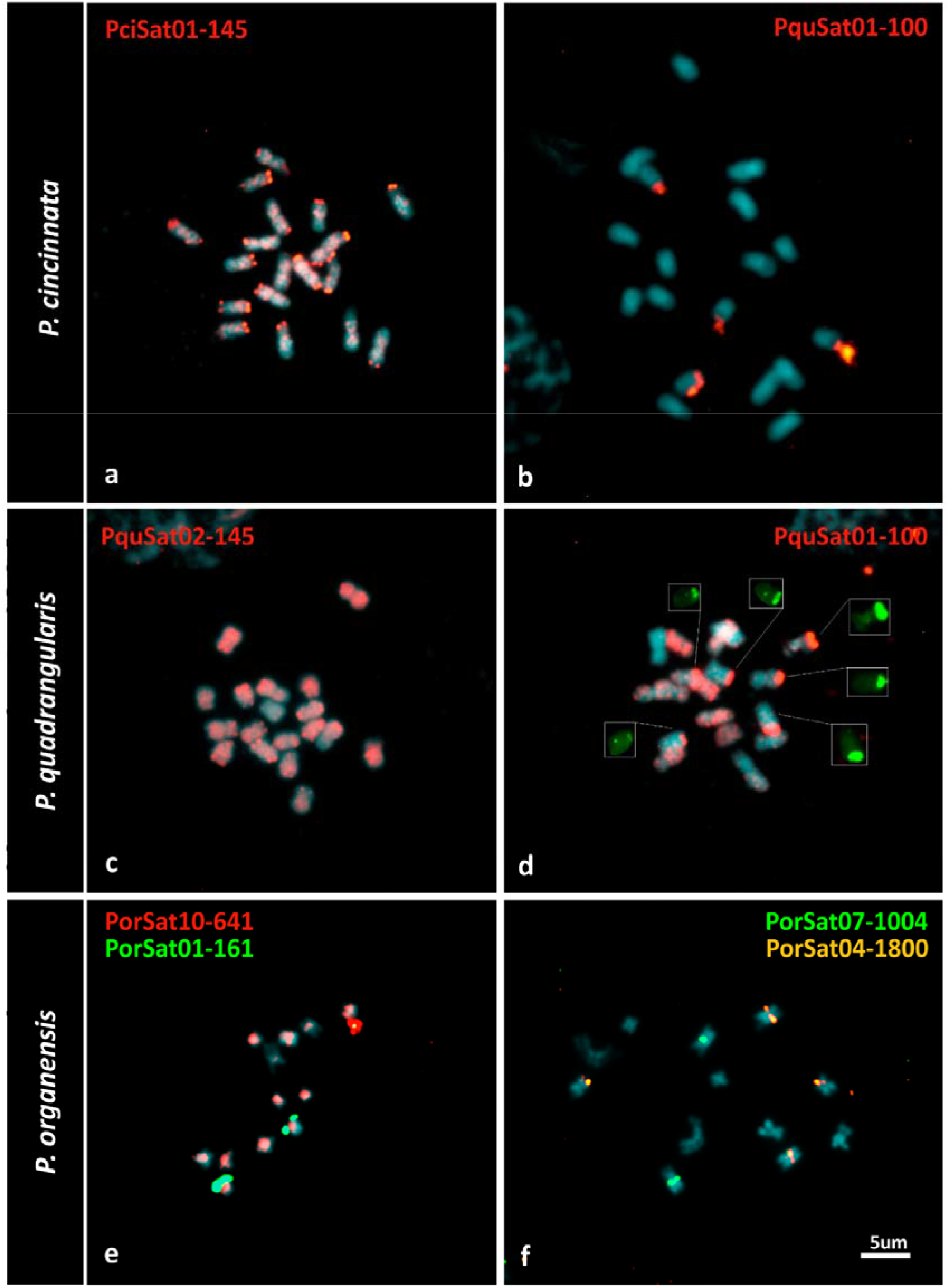
Localization of different satellite DNA repeats (a-f) and 35S rDNA (d) on mitotic chromosomes of *Passiflora* species. (**a-b**) *Passiflora cincinnata*, (**c-d**) *P. quadrangularis*, and (**e-f**) *P. organensis*. Hybridization signals are in the same colour of repeat names in each picture.

In *P. organensis*, PorSat01-161 showed a pair of subterminal signals in one and the same chromosome pair (Fig. 5e). PorSat04-1800 showed two pair of sites, a subterminal and a pericentromeric, in different chromosome pairs (Fig. 5f). PorSat07-1004 showed pericentromeric signals in another chromosome pair (Fig. 5f). These four repeats are useful markers for chromosome identification. PorSat10-641 showed mainly pericentromeric distribution in all chromosome pairs, with variable intensities (Fig. 5a). PorSat05-510 showed dispersed distribution in all chromosome pairs, while PorSat12-52, PorSat02-371 and PorSat22-398 showed a scattered distribution throughout the genome (data not shown).

### Relations between satellites and the 35S rDNA

Because PquSat01-100 signals were co-localized with the 35S rDNA sites, we searched for similarities between these repeats. After assembling the *Passiflora* 35S rDNA, we have performed automatic gene annotation and described the gene sequences corresponding to the 26S, 18S and 5.8S genes and the transcribed spacer regions (ITS1 and ITS2) (Online Resource 2). Also, using bioinformatics tools we have observed high similarity (100% identity) between PquSat01-100 satellite and the IGS region of the 35S rDNA sequence. The region of similarity is made up of four subunits, two of 595 and two of 344, with smaller (100 to 200bp) subrepeats (Online Resource 3). We hypothesize that PquSat01-100 satellite derives from part of IGS, from which it amplified and dispersed in the *P. quadrangularis* genome.

## DISCUSSION

In the present work, we report the first comparative study of repetitive elements using genomic *in silico* analysis and cytogenomics for comprehensive characterization of the tenfold genome size variation in *Passiflora* (Yotoko et al. 2011; Souza et al. 2004). Plant genome sizes span several orders of magnitude ranging from the 63–64 Mbp in *Genlisea* spp. (Fleischmann et al. 2014) to the more than 148,851 Mbp genome of *Paris japonica* (Pellicer et al. 2010). Significant genome size variations are also present within other genera, such as *Gossypium, Oryza* and *Cuscuta*, for which 3, 3.6 and 102-fold genome size variations, respectively (Ammiraju et al. 2006; Hendrix and Stewart, 2005; Neumann et al. 2020), but the variation observed in *Passiflora* is among the highest for a single genus in angiosperms. This genome size variation in plants is mainly due to polyploidization (Adams and Wendel, 2005; Bennetzen et al. 2005) and TE proliferation and/or elimination (Devos et al. 2002; Hawkins et al. 2009; Ma et al. 2004; Neumann et al. 2011). We observed that about 28 to 73% of the genome is composed by TEs and between 0,1% and 4% by satellite DNA, which is comparable to other plant genomes of similar sizes (Macas et al. 2011; Macas et al. 2015). Such differences in percentage of TE explains most of the difference in genome size in *Passiflora*, with larger genomes having correspondingly higher amounts of repeats.

As in most angiosperms (Weiss-Schneeweiss et al. 2015), Ty3/gypsy dominated the repetitive fraction of *P. quadrangularis* genome (35%), represented mainly by the lineages Tekay and Athila. However, Ty1/copia-Angela was highly abundant with 24%. In contrast, Ty1/copia retrotransposons showed higher proportion (35.7%) in *P. cincinnata*, due to an even higher proportion of Angela (29%). This high abundance of Ty1/copia has already been observed in the passion-fruit, *Passiflora edulis* (16.89% versus 33.33% for Ty3/gypsy, Pamponét et al. 2019), another species from the *Passiflora* subgenus but more closely related to *P. quadrangularis* than to *P. cincinatta* (Sader et al 2019a). In *Passiflora edulis* (1,232 Mpb, Yotoko et al. 2011), Costa et al. (2019) corroborated Angela as the most abundant lineage (1.9%) from the Ty1/copia superfamily (3.12%), although -Tekay (8.5%, referred to as *Del*) from Ty3/gypsy (10.52%) was more abundant in their analysis. In *P. organensis*, Tekay accounted for most of its repetitive fraction as seen in *P. quadrangularis*, Angela was not detected, suggesting it might have played a significant role in its reduced genome size. Thus, the increase in genome size within the genus, so far apparently concentrated in the *Passiflora* subgenus, was caused by independent patterns of expansions of both Ty3/gypsy, with 33% in *P. edulis* (Pamponét et al. 2019), 18.45% in *P. cincinnata* and 41.22% in *P. quadrangularis*, and Ty1/copia, with 35.66% in *P. cincinnata*, 25.44% in *P. quadrangularis* and 16.89% in *P. edulis*.

We observed that most transposable element families are represented only by two or three clusters indicating their long-term presence without changes in sequence or structure. Just Athila, Angela and Tekay (and SIRE in *P. cincinnata* only) retrotransposons were found in multiple clusters suggesting higher divergence and abundance. Most of the Tekay elements were incomplete or truncated in the three species, suggesting active elimination. Thus, the increase in genome size may be due to large and long-term bursts, so far not counterbalanced by sufficient removal mechanisms (Ibarra-Laclette et al. 2013). The opposite was observed in *Fritillaria*; where the evolution of truly obese genomes was largely determined by the failure of the mechanisms responsible for repeated elimination that effectively operate in species with smaller genomes to counteract genome expansion (Ambrozová et al. 2011). Transposable elements are frequently recognized as “genomic fossils” that were once autonomous, but, at some point, they experience mutations that leave them inactive (Cruz et al. 2014). The majority of *P. edulis* TEs (70.8%) were incomplete, corroborating previous findings showing that most TE copies are either defective or fossilized (Costa et al. 2019). Only Angela showed a higher proportion of complete elements in *P. edulis* (Costa et al. 2019), compatible with a burst of amplification restricted to the subgenus *Passiflora*.

Divergence in repetitive DNA is a primary driving force for genome and chromosome evolution. The relationships of full-length Tekay elements from the three species in the phylogenetic reconstruction showed that most of the clusters are shared by *P. cincinnata* and *P. quadrangularis*. This similarity probably reflects that they already existed 12 Mya in the common ancestor of the clade that contains these two species (Sader et al. 2019a). The variants of the clusters that are shared by the three species belong to a Tekay that probably was already present in the common ancestor of the two subgenera, and before its evolutionary divergence >40 Mya (Sader et al. 2019a). According to Costa et al. (2019), insertions of retroelements in *P. edulis* were dated between only one and two Mya. In our study, we have seen that some Tekay clusters were present in the last common ancestral among the three species. Apart from methodological differences (we are investigating clusters of TEs, not individual elements), a possible explanation for this discrepancy could be that the authors analysed TEs from gene-rich regions, which are located at chromosomal ends (Sader et al. 2019b). In our case, we used elements from the most abundant clusters that may have accumulated in the pericentromeric regions for longer time.

*In situ* localization of the retrotransposons in pericentromeric regions or dispersed throughout the chromosomes is a common feature of plant genomes of similar sizes, small and large, respectively (Miller et al. 1998; Cheng and Murata 2003; Nagaki et al. 2004, Neumann et al. 2011). Unlike retrotransposons, satellite DNAs usually form blocks on heterochromatic chromosome regions (Heslop-Harrison and Schwarzacher 2011; Heslop-Harrison and Schmidt 2012; Ribeiro et al. 2017). Here we have observed that Ty1/copia-Angela and Ty3/gypsy-Tekay retroelements were dispersed in *P. quadrangularis* and *P. cincinnata* chromosomes, although showing more intense labelling at proximal regions. An uneven dispersed distribution was also observed in *P. edulis* and two other species of the subgenus *Passiflora*, where terminal or subterminal regions of the chromosome arms are gene-rich, and the proximal regions are gene-poor and consist of dispersed repetitive sequences (Pamponét et al. 2019; Sader et al. 2019b; Stack et al. 2009; Dias et al. 2020). This distribution pattern suggest that gene-rich regions, and probably recombination, is more evenly distributed in *P. organensis*, except in pericentromeric regions, while higher in chromosomal ends of larger genome species, such as *P. quadrangularis, P. cincinnata*, and *P. edulis*.

Contrasting to transposable elements, differences in abundance of satellite DNAs showed that these repeats have not contributed to DNA content differences in the genus. *Passiflora organensis*, the species with the smaller genome, contains the largest proportion (1.5-4%) and an unusually large number of different satDNA. This wide diversity in such a small genome size with 46 satDNA different families is not commonly observed in plants, only in *Luzula* (Heckman et al. 2013) and in *Vicia faba* (Robledillo et al. 2018). Different satDNA families may be present in one species. For example, there are up to 15 families in *Pisum sativum* (Macas et al. 2007), 62 families in *Locusta migratoria* (Ruiz-Ruano et al. 2016), or nine satDNA families within the human genome (Levy et al. 2007; Miga 2015). However, there are usually one or a few predominant satDNA families in each species (Macas et al. 2007; Ruiz-Ruano et al. 2016; Levy et al. 2007; Miga 2015). In *Passiflora*, we have observed only one superfamily (SF1) of satDNA, with a subtelomeric distribution, shared by all three species studied here and in *P. edulis* (Pamponét et al. 2019). The other repeats were mostly species-specific, suggesting that there are different amplification and diversification patterns for this repetitive fraction in the genus. This low degree of sharing of most satDNA families is possibly related to the long divergence time between subgenus *Passiflora* and *Decaloba* genomes and among species of the same subgenus (Sader et al. 2019a).

With the aim of confirming the low degree of satDNA sharing observed after RepeatExplorer comparative analysis, we used bioinformatics tools to search for all identified satDNAs in the other two species. Only one of these satDNA clusters (PorSat20-3100) was also detected in both species of the *Passiflora* subgenus (although in very low proportions) and PorSat37-970 was also found in *P. quadrangularis*. This can be due to the fact that, in larger genomes, satDNAs present in low or very low abundance failed to be detected using RepeatExplorer pipeline. Using satMiner, the search for satDNA was more efficient. Nevertheless, satDNAs which occur in very low abundance, such as PorSat20-3100 and PorSat37-970, were only possible to find using the Map to reference tool in Geneious. Thus, for large plant genomes, where it may be difficult to find satellites in low abundance, we suggest the use of the three combined approaches. Thus, the comparative analyses with species with smaller genomes may be a good option for finding satDNA in larger genomes using this approach.

Satellite repeats may occur at subtelomeric or instersticial chromosome regions, but preferentially in centromeres (Garrido-Ramos 2015). We have found one subtelomeric superfamily (SF1) of satDNA present in all three species studied here and in *P. edulis* (Pamponét et al. 2019), but no conserved putative centromeric repeat was found. Conserved centromeric repeats in a genus or beyond is rare (Zhong et al. 2002), but conserved subtelomeric repeat have been previously found for example in *Phaseolus* (Ribeiro et al. 2019). Furthermore, the CRM elements, which are centromeric in other species such as maize (Zhong et al. 2002), are in very low abundance in *Passiflora* (0.32% in *P. quadrangularis;* 0.37% in *P. cincinnata;* and 0.31% in *P. organensis)*. Therefore, the nature of *Passiflora* centromeres needs to be further investigated. The PquSat01-100, observed in *P. quadrangularis* and *P. cincinnata*, was probably originated from the IGS regions of the 35S rDNA. There are several examples where satDNA originated from the IGS or rDNA coding genes (Garrido-Ramos 2015; Plohl 2012; Kirov et al. 2018) for example, the satDNA *jumper*, in the *Phaseolus* genus, derives from the NTS of the 5S rDNA (Ribeiro et al. 2017), showing that this phenomenon is quite common, even in plants in low proportions of satDNA and TEs increased genomes.

## CONCLUSIONS

This is the first comparative study of the repetitive fraction in the *Passiflora* genus, expanding the extremes of genomic sizes, and including two subgenus (*Passiflora* and *Decaloba*) and further two cultivated species (*P. cincinnata* and *P. quadrangularis*). Our results showed that *P. quadrangularis* presents a higher accumulation of repetitive DNA sequences, but less divergence in relation to *P. organensis*. *Passiflora cincinnata* showed similarity to *P. quadrangularis* regarding the families of repetitive DNA sequences, although in different proportions, probably reflecting phylogenetic relationships. *Passiflora organensis* presented greater diversity and the highest proportion of satDNA. Together, our data pointed out that the satellitome is not the fraction responsible for the increase in genome size in *Passiflora*. This increase was originated by the expansion, of two main retroelement lineages, (Ty3/gypsy-Tekay and Ty1/copia-Angela retrotransposons.

## Supporting information

Online Resource

## ABBREVIATIONS

Cy3-dUTP: 5-amino-propargyl-2’-deoxyuridine 5’-triphosphate coupled to red cyanine fluorescent dye
DAPI: 4’, 6-diamidino-2-phenylindole
FISH: Fluorescent *in situ* hybridization
LTR: Long terminal repeat
NGS: Next-generation sequencing
rDNA: Ribossomal DNA
RT: Retrotransposons
satDNA: Satellite DNA
TAREAN: TAndem REpeat ANalyser
TEs: Transposable elements

## Compliance with ethical standards

Conflict of interest: The authors declare that they have no conflict of interest.

## Author contribution statement

MAS designed experiments, performed bioinformatic analysis, amplifications, probe labeling, FISH, data analysis and wrote the manuscript. MV analysed data and assisted with writing the manuscript; LAC, MLC, NFM and MCD provided plant material and whole-genome sequencing. APH designed the experiments, analysed data and wrote the manuscript. All authors read and approved the manuscript.

## Acknowledgements

We are grateful to Dr. Francisco Ruiz-Ruano for bioinformatics support and for critical reading of the manuscript. This study was partially supported by Fundação de Amparo à Pesquisa do Estado de Pernambuco (FACEPE) through scholarships awarded to MAS (IBPG-1086-2.03/15 and FACEPE AMD-0128-2.00/17) and CNPq (Conselho Nacional de Desenvolvimento Científico e Tecnológico) through fellowship awarded to AP-H. This study was supported in part by the Coordenacão de Aperfeiçoamento de Pessoal de Nivel Superior - Brasil (CAPES, Finance Code 001); EMBRAPA (Embrapa SEG-02.16.04.007.00.03) and FAPESP (2019/07838-6).

## Notes

### Competing Interest Statement

The authors have declared no competing interest.

## References

Adams KL, Wendel JF (2005) Polyploidy and genome evolution in plants. Current Opinion on Plant Biology 8:135–141. https://doi.org/10.1016/j.pbi.2005.01.001

Albach DC & Greilhuber J (2004) Genome size variation and evolution in Veronica. Annals of Botany 94:897–911. https://doi.org/10.1093/aob/mch219

Ambrožová K, Mandáková T, Bureš P, Neumann P, Leitch IJ, Koblížková A, … & Lysak MA (2011) Diverse retrotransposon families and an AT-rich satellite DNA revealed in giant genomes of *Fritillaria* lilies. Annals of Botany 107:255–268. https://doi.org/10.1093/aob/mcq235

Ammiraju JS, Luo M, Goicoechea JL, Wang W, Kudrna D, Mueller C & Fang E (2006) The *Oryza* bacterial artificial chromosome library resource: construction and analysis of 12 deep-coverage large-insert BAC libraries that represent the 10-genome types of the genus *Oryza*. Genome Research 16:140–147. http://www.genome.org/cgi/doi/10.1101/gr.3766306.

Araya S, Martins AM, Junqueira NT, Costa AM, Faleiro FG & Ferreira ME (2017) Microsatellite marker development by partial sequencing of the sour passion fruit genome *(Passiflora edulis* Sims). BMC Genomics 18:549. https://doi.org/10.1186/s12864-017-3881-5

Aversano R, Contaldi F, Ercolano MR, Grosso V, Iorizzo M, Tatino F & Delledonne M (2015) The *Solanum commersonii* genome sequence provides insights into adaptation to stress conditions and genome evolution of wild potato relatives. The Plant Cell 27:954–968. https://doi.org/10.1105/tpc.114.135954

Bennetzen JL, Ma J, Devos KM (2005) Mechanisms of recent genome size variation in flowering plants. Annals of Botany 95:127–132. https://doi.org/10.1093/aob/mci008

Biscotti MA, Olmo E & Heslop-Harrison JP (2015) Repetitive DNA in eukaryotic genomes. 415–420. https://doi.org/10.1007/s10577-015-9499-z

Carvalho CR, & Saraiva LS (1993) An air-drying technique for maize chromosomes without enzymatic maceration. Biotechnic & Histochemistry 68:142–145. https://doi.org/10.3109/10520299309104684

Cheng ZJ & Murata M (2003) A centromeric tandem repeat family originating from a part of Ty3/gypsy-retroelement in wheat and its relatives. Genetics 164:665–672.

Costa ZP, Cauz-Santos LA, Ragagnin GT, Van Sluys MA, Dornelas MC, Berges H & Vieira MLC (2019) Transposable element discovery and characterization of LTR-retrotransposon evolutionary lineages in the tropical fruit species *Passiflora edulis*. Molecular Biology Reports 46:6117–6133. https://doi.org/10.1007/s11033-019-05047-4

Cruz GMQ, Metcalfe CJ, De Setta N et al (2014) Virus-like attachment sites and plastic CpG Islands: landmarks of diversity in plant Del retrotransposons. PLoS ONE 9:1–14. https://doi.org/10.1371/journal.pone.0097099

De Koning APJ, Gu W, Castoe TA, Batzer MA, Pollock DD (2011) Repetitive elements may comprise over two-thirds of the human genome. PLoS Genetics 7:e1002384. https://doi:10.1371/journal.pgen.1002384

Derks MF, Smit S, Salis L, Schijlen E, Bossers A, Mateman C & Megens HJ (2015) The genome of winter moth (*Operophtera brumata*) provides a genomic perspective on sexual dimorphism and phenology. Genome Biology and Evolution 7:2321–2332. https://doi.org/10.1093/gbe/evv145

Devos KM, Brown JK & Bennetzen JL (2002) Genome size reduction through illegitimate recombination counteracts genome expansion in *Arabidopsis*. Genome Research 12:1075–1079. https://doi.org/10.1101/gr.132102

Dias Y, Sader MA, Vieira ML, & Pedrosa-Harand A (2020) Comparative cytogenetic maps of *Passiflora alata* and *P. watsoniana* (Passifloraceae) using BAC-FISH. Plant Systematics and Evolution 306:1–8. https://doi.org/10.1007/s00606-020-01675-7

Dierckxsens N, Mardulyn P & Smits G (2016) NOVOPlasty: de novo assembly of organelle genomes from whole genome data. Nucleic Acids Research 45:e18–e18. https://doi.org/10.1093/nar/gkw955

Fleischmann A, Michael TP, Rivadavia F, Sousa A, Wang W, Temsch EM & Heubl G (2014) Evolution of genome size and chromosome number in the carnivorous plant genus *Genlisea* (Lentibulariaceae), with a new estimate of the minimum genome size in angiosperms. Annals of Botany 114:1651–1663. https://doi.org/10.1093/aob/mcu189

Fonsêca A, Ferreira J, Dos Santos TRB, Mosiolek M, Bellucci E, Kami J & Pedrosa-Harand A (2010) Cytogenetic map of common bean *(Phaseolus vulgaris* L.) Chromosome Research 18:487–502. https://doi.org/10.1007/s10577-010-9129-8

Gaiero P, Vaio M, Peters SA, Schranz ME, de Jong H, & Speranza PR (2019) Comparative analysis of repetitive sequences among species from the potato and the tomato clades. Annals of Botany 123:521–532. https://doi.org/10.1093/aob/mcy186

Garrido-Ramos MA (2015) Satellite DNA in plants: More than Just Rubbish. Genomics and Informatics 12:87. https://doi.org/10.1159/000437008

Garrido-Ramos MA (2017) Satellite DNA: An evolving topic. Genes 9:230. https://doi.org/10.3390/genes8090230

Gerlach WL, & Bedbrook JR (1979) Cloning and characterization of ribosomal RNA genes from wheat and barley. Nucleic Acids Research 7:1869–1885. https://doi.org/10.1093/nar/7.7.1869

Goubert C, Modolo L, Vieira C, Valiente Moro C, Mavingui P & Boulesteix M (2015) De novo assembly and annotation of the Asian tiger mosquito *(Aedes albopictus)* repeatome with dnaPipeTE from raw genomic reads and comparative analysis with the yellow fever mosquito (*Aedes aegypti*). Genome Biology and Evolution 7:1192–1205. https://doi.org/10.1093/gbe/evv050

Hannan AJ (2018) Tandem Repeats and Repeatomes: Delving Deeper into the ‘Dark Matter’of Genomes. EBioMedicine 31:3–4. https://doi.org/10.1016/j.ebiom.2018.04.004

Hansen AK, Gilbert LE, Simpson BB, Downie SR, Cervi AC, and Jansen RK (2006) Phylogenetic relationships and chromosome number evolution in *Passiflora*. Systematic Botany 31:138–150. https://doi.org/10.1600/036364406775971769

Hawkins JS, Proulx SR, Rapp RA, Wendel JF (2009) Rapid DNA loss as a counterbalance to genome expansion through retrotransposon proliferation in plants. Proceedings of the National Academy of Sciences USA 106:17811–17816. https://doi.org/10.1073/pnas.0904339106

Heckmann S, Macas J, Kumke K, Fuchs J, Schubert V, Ma L, … & Houben A (2013) The holocentric species *Luzula elegans* shows interplay between centromere and large scale genome organization. The Plant Journal, 73:555–565. https://doi.org/10.1111/tpj.12054

Hemleben V, Kovarik A, Torres□Ruiz RA, Volkov RA, & Beridze T (2007) Plant highly repeated satellite DNA: molecular evolution, distribution and use for identification of hybrids. Systematics and Biodiversity 5:277–289. https://doi.org/10.1017/S147720000700240X

Hendrix B, & Stewart JM (2005) Estimation of the nuclear DNA content of *Gossypium* species. Annals of Botany 95:789–797. https://doi.org/10.1093/aob/mci078

Heslop-Harrison JSP, & Schwarzacher T (2011) Organisation of the plant genome in chromosomes. The Plant Journal 66:18–33. https://doi.org/10.1111/j.1365-313X.2011.04544.x

Heslop□Harrison JP, & Schmidt T (2012) Plant nuclear genome composition. eLS. https://doi.org/10.1002/9780470015902.a0002014.pub2

Ibarra-Laclette E, Lyons E, Hernández-Guzmán G, et al (2013) Architecture and evolution of a minute plant genome. Nature 498:94–98. https://doi.org/10.1038/nature12132

Jouffroy O, Saha S, Mueller L, Quesneville H & Maumus F (2016) Comprehensive repeatome annotation reveals strong potential impact of repetitive elements on tomato ripening. BMC Genomics 17:624. https://doi.org/10.1186/s12864-016-2980-z

Judd WS, Campbell CC, Kellogg EA, Stevens PF & Donoghue MJ (2015) Plant systematics: A phylogenetic approach 3^rd^ ed. Sinauer Associates Inc. Sunderland Massachusetts

Katoh K & Standley DM (2013) MAFFT multiple sequence alignment software version 7: improvements in performance and usability. Molecular Biology and Evolution 30:772–780. https://doi.org/10.1093/molbev/mst010

Kearse M, Moir R, Wilson A, Stones-Havas S, Cheung M, Sturrock S & Thierer T (2012) Geneious Basic: an integrated and extendable desktop software platform for the organization and analysis of sequence data. Bioinformatics 28:1647–1649. https://doi.org/10.1093/bioinformatics/bts199

Kidwell Margaret G (2002) Transposable elements and the evolution of genome size in eukaryotes. Genetica 115:49–63. https://doi.org/10.1023/A:1016072014259

King K, Jobst J & Hemleben V (1995) Differential homogenization and amplification of two satellite DNAs in the genus *Cucurbita* (Cucurbitaceae). Journal of Molecular Evolution 41:996–1005. https://doi.org/10.1007/BF00173181

Kirov I, Gilyok M, Knyazev A, & Fesenko I (2018) Pilot satellitome analysis of the model plant, *Physcomitrella patens*, revealed a transcribed and high-copy IGS related tandem repeat. Comparative Cytogenetics 12:493. https://doi.org/10.3897/CompCytogen.v12i4.31015

Lander ES, Linton LM, Birren B, Nusbaum C, Zody MC Baldwin J, Devon K, Dewar K, Doyle M, FitzHugh W et al. (2001) Initial sequencing and analysis of the human genome. Nature 409:860–921. https://doi.org/10.1038/35087627

Levin HL & Moran JV (2011) Dynamic interactions between transposable elements and their hosts. Nature Reviews Genetics 12:615. https://doi.org/10.1038/nrg3030

Levy S, Sutton G, Ng PC, Feuk L, Halpern AL, Walenz BP, Axelrod N, Huang J, Kirkness EF, Denisov G, et al (2007) The diploid genome sequence of an individual human. PLoS Biology 5:e254. https://doi.org/10.1371/journal.pbio.0050254

López-Flores I & Garrido-Ramos MA (2012) The repetitive DNA content of eukaryotic genomes. In Repetitive DNA 7:1–28. Karger Publishers. https://doi.org/10.1159/000337118

Ma J, Devos KM, Bennetzen JL (2004) Analyses of LTR-retrotransposon structures reveal recent and rapid genomic DNA loss in rice. Genome Research 14:860–869. http://www.genome.org/cgi/doi/10.1101/gr.1466204

Macas J, Neumann P & Navrátilová A (2007) Repetitive DNA in the pea *(Pisum sativum* L.) genome: comprehensive characterization using 454 sequencing and comparison to soybean and *Medicago truncatula*. BMC Genomics 8:427. https://doi.org/10.1186/1471-2164-8-427

Macas J, Kejnovský E, Neumann P, Novák P, Koblížková A, Vyskot B (2011) Next generation sequencing-based analysis of repetitive DNA in the model Dioceous plant *Silene latifolia*. PLoS One 6:11. https://doi.org/10.1371/journal.pone.0027335

Macas J, Novak P, Pellicer J, Čížková J, Koblížková A, Neumann P & Leitch IJ (2015) In depth characterization of repetitive DNA in 23 plant genomes reveals sources of genome size variation in the legume tribe Fabeae. PLoS One 10:11. https://doi.org/10.1371/journal.pone.0143424

Margulies M, Egholm M, Altman WE, Attiya S, Bader JS, Bemben LA & Dewell SB (2005) Genome sequencing in microfabricated high-density picolitre reactors. Nature 437:376. https://doi.org/10.1038/nature03959

Maumus F, Quesneville H (2016) Impact and insights from ancient repetitive elements in plant genomes. Current Opinion in Plant Biology 30:41–6. https://doi.org/10.1016/j.pbi.2016.01.003

McCann J, Macas J, Novák P, Stuessy TF, Villaseñor JL, & Weiss-Schneeweiss H (2020) Differential Genome Size and Repetitive DNA Evolution in Diploid Species of *Melampodium* sect. Melampodium (Asteraceae). Frontiers in Plant Science 11:362. https://doi.org/10.3389/fpls.2020.00362

Melo NF, Guerra FMS (2003) Variability of the 5S and 45SrDNA sites in *Passiflora* L. species with distinct base chromosome numbers. Annals of Botany 92:309–316. https://doi.org/10.1093/aob/mcg138

Metzlaff M, Troebner W, Baldauf F, Schlegel R & Cullum J (1986) Wheat specific repetitive DNA sequences—construction and characterization of four different genomic clones. Theoretical and Applied Genetics 72:207–210.

Miga KH (2015) Completing the human genome: The progress and challenge of satellite DNA assembly. Chromosome Research. 23:421–426. https://doi.org/10.1007/s10577-015-9488-2

Mikkelsen TS, Wakefield MJ Aken B, Amemiya CT Chang JL, Duke S, Garber M, Gentles AJ, Goodstadt L, Heger A et al (2007) Genome of the marsupial *Monodelphis domestica* reveals innovation in non-coding sequences. Nature 447:167–178. https://doi.org/doi:10.1038/nature05805

Miller JT, Dong F, Jackson SA, Song J & Jiang J (1998) Retrotransposon-related DNA sequences in the centromeres of grass chromosomes. Genetics 150:1615–1623.

Nagaki K et al (2004) Sequencing of a rice centromere uncovers active genes. Nature genetics 36:138. https://doi.org/doi:10.1038/ng1289

Neumann P et al (2020) Impact of parasitic lifestyle and different types of centromere organization on chromosome and genome evolution in the plant genus *Cuscuta*. bioRxiv. https://doi.org/10.1101/2020.07.03.186437

Neumann P, Novák P, Hoštáková N & Macas J (2019) Systematic survey of plant LTR-retrotransposons elucidates phylogenetic relationships of their polyprotein domains and provides a reference for element classification. Mobile DNA 10:1. https://doi.org/10.1186/s13100-018-0144-1

Neumann P, Navrátilová A, Koblížková A, Kejnovský E, Hřibová E, Hobza R & Macas J (2011) Plant centromeric retrotransposons: a structural and cytogenetic perspective. Mobile DNA 2:4. http://www.mobilednajournal.com/content/2/1/4

Novák P, Neumann P & Macas J (2010) Graph-based clustering and characterization of repetitive sequences in next-generation sequencing data. BMC Bioinformatics 11:378. http://www.biomedcentral.com/1471-2105/11/378

Novák P, Neumann P, Pech J, Steinhaisl J & Macas J (2013) RepeatExplorer: a Galaxy-based web server for genome-wide characterization of eukaryotic repetitive elements from nextgeneration sequence reads. Bioinformatics 29:792–793. https://doi.org/10.1093/bioinformatics/btt054

Novák P, Ávila LR, Koblížková A, Vrbová I, Neumann P, Macas J (2017) TAREAN: a computational tool for identification and characterization of satellite DNA from unassembled shortreads. Nucleic Acids Research 45:e111. https://doi.org/10.1093/nar/gkx257

Pamponét VCC et al (2019) Correction to: low coverage sequencing for repetitive DNA analysis in *Passiflora edulis* Sims: citogenomic characterization of transposable elements and satellite DNA. BMC Genomics 20:303. https://doi.org/10.1186/s12864-019-5576-6

Pellicer J, Fay MF, Leitch IJ (2010) The largest eukaryotic genome of them all? Botanical Journal of the Linnean Society 164:10–15. https://doi.org/10.1111/j.1095-8339.2010.01072.x

Pita S, Panzera F, Mora P, Vela J, Cuadrado Á, Sánchez A & Lorite P (2017) Comparative repeatome analysis on *Triatoma infestans* Andean and Non-Andean lineages, main vector of Chagas disease. PLoS One 12:e0181635. https://doi.org/10.1371/ioumal.pone.0181635

Plohl M, Meštrović N, Mravinac B (2012) Satellite DNA evolution. In Repetitive DNA 7:126–152. Karger Publishers. https://doi.org/10.1159/000337122

Pritham EJ (2009) Transposable elements and factors influencing their success in eukaryotes. Journal of Heredity 100:648–655. https://doi.org/10.1093/jhered/esp065

Rayburn AL, & Gill BS (1986) Isolation of a D-genome specific repeated DNA sequence from *Aegilops squarrosa*. Plant Molecular Biology Reporter 4:102–109. https://doi.org/10.1007/BF02732107

Ribeiro T, Vasconcelos E, Dos Santos KG, Vaio M, Brasileiro-Vidal AC & Pedrosa-Harand A (2019) Diversity of repetitive sequences within compact genomes of *Phaseolus* L. beans and allied genera *Cajanus* L. and *Vigna* Savi. Chromosome Research 1–15. https://doi.org/10.1007/s10577-019-09618-w

Ribeiro T, Dos Santos KG, Richard MM, Sévignac M, Thareau V, Geffroy V & Pedrosa-Harand A (2017) Evolutionary dynamics of satellite DNA repeats from *Phaseolus* beans. Protoplasma 254:791–801. https://doi.org/10.1007/s00709-016-0993-8

Robledillo LÁ, Koblížková A, Novák P, Böttinger K, Vrbová I, Neumann P, … & Macas J (2018) Satellite DNA in *Vicia faba* is characterized by remarkable diversity in its sequence composition, association with centromeres, and replication timing. Scientific Reports 8:1–11. https://doi.org/10.1038/s41598-018-24196-3

Ruiz-Ruano FJ, López-León MD, Cabrero J & Camacho JPM (2016) High-throughput analysis of the satellitome illuminates satellite DNA evolution. Scientific Reports 6:28333. https://doi.org/10.1038/srep28333

Sader MA, Amorim BS, Costa L, Souza G & Pedrosa-Harand A (2019a) The role of chromosome changes in the diversification of *Passiflora* L. (Passifloraceae). Systematics and Biodiversity 17:7–21. https://doi.org/10.1080/14772000.2018.1546777

Sader MA, Dias Y, Costa ZP, Munhoz C, Penha H, Bergès H & Pedrosa-Harand A (2019b) Identification of passion fruit *(Passiflora edulis)* chromosomes using BAC-FISH. Chromosome Research 1–13. https://doi.org/10.1007/s10577-020-09633-2

Schnable PS, Ware D, Fulton RS, Stein JC, Wei F, Pasternak S, Liang C, Zhang J, Fulton L, Graves TA et al (2009) The B73 maize genome: Complexity, diversity, and dynamics. Science 326:1112–1115. https://doi.org/10.1126/science.1178534

Siefert, J. L. (2009). Defining the mobilome. In Horizontal gene transfer (pp. 13–27). Humana Press. https://doi.org/10.1007/978-1-60327-853-9_2

Souza MM, Palomino G, Pereira MG, Viana AP (2004) Flow cytometric analysis of genome size variation in some *Passiflora* species. Hereditas 141:31–38. https://doi.org/10.1111/j.1601-5223.2004.01739.x

Stack SM, Royer SM, Shearer LA, Chang SB, Giovannoni JJ, Westfall DH & Anderson LK (2009) Role of fluorescence *in situ* hybridization in sequencing the tomato genome. Cytogenetic and Genome Research 124:339–350. https://doi.org/10.1159/000218137

Ulmer T & Macdougal JM (2004) *Passiflora:* Passion flowers of the World. Timber Press, Portland.

Untergasser A, Cutcutache I, Koressaar T, Ye J, Faircloth BC, Remm M & Rozen SG (2012). Primer3–new capabilities and interfaces. Nucleic Acids Research. https://doi.org/10.1093/nar/gks596

Van-Lume, B., Mata-Sucre, Y., Báez, M., Ribeiro, T., Huettel, B., Gagnon, E., … & Souza, G. (2019). Evolutionary convergence or homology? Comparative cytogenomics of Caesalpinia group species (Leguminosae) reveals diversification in the pericentromeric heterochromatic composition. Planta 250:2173–2186. http://dx.doi.org/10.1007/s00425-019-03287-z

Weising K, Nybom H, Pfenninger M, Wolff K & Kahl G (2005) DNA fingerprinting in plants: principles, methods, and applications. CRC press.

Weiss-Schneeweiss H, Leitch AR, McCann J, Jang TS, & Macas J (2015) Employing next generation sequencing to explore the repeat landscape of the plant genome. Next generation sequencing in plant systematics. Regnum Vegetabile 157:155–179. http://dx.doi.org/10.14630/000006

Wicker T, Sabot F, Hua-Van A, Bennetzen JL, Capy P, Chalhoub B & Paux E (2007) A unified classification system for eukaryotic transposable elements. Nature Reviews Genetics 8:973–982. https://doi.org/10.1038/nrg2165-c4

Wolf PG, Sessa EB, Marchant DB, Li FW, Rothfels CJ, Sigel EM & Soltis PS (2015) An exploration into fern genome space. Genome Biology and Evolution 7:2533–2544. https://doi.org/10.1093/gbe/evv163

Yotoko KSC, Dornelas MC, Togni PD, Fonsêca TC, Salzano FM, Bonatto SL, Freitas LB (2011) Does variation in genome sizes reflect adaptive or neutral processes? New clues from *Passiflora*. Plos One 6:127–131. https://doi.org/10.1371/journal.pone.0018212

Zhong CX, Marshall JB, Topo C, Mroczek R, Kato A, Nagaki K & Dawe RK (2002) Centromeric retroelements and satDNA interact with maize kinetochore protein CENH3. The Plant Cell 14:2825–2836. https://doi.org/10.1105/tpc.006106

