## Supplementary material for "Large vs small genomes in *Passiflora*: the influence of the mobilome and the satellitome": Online Resource

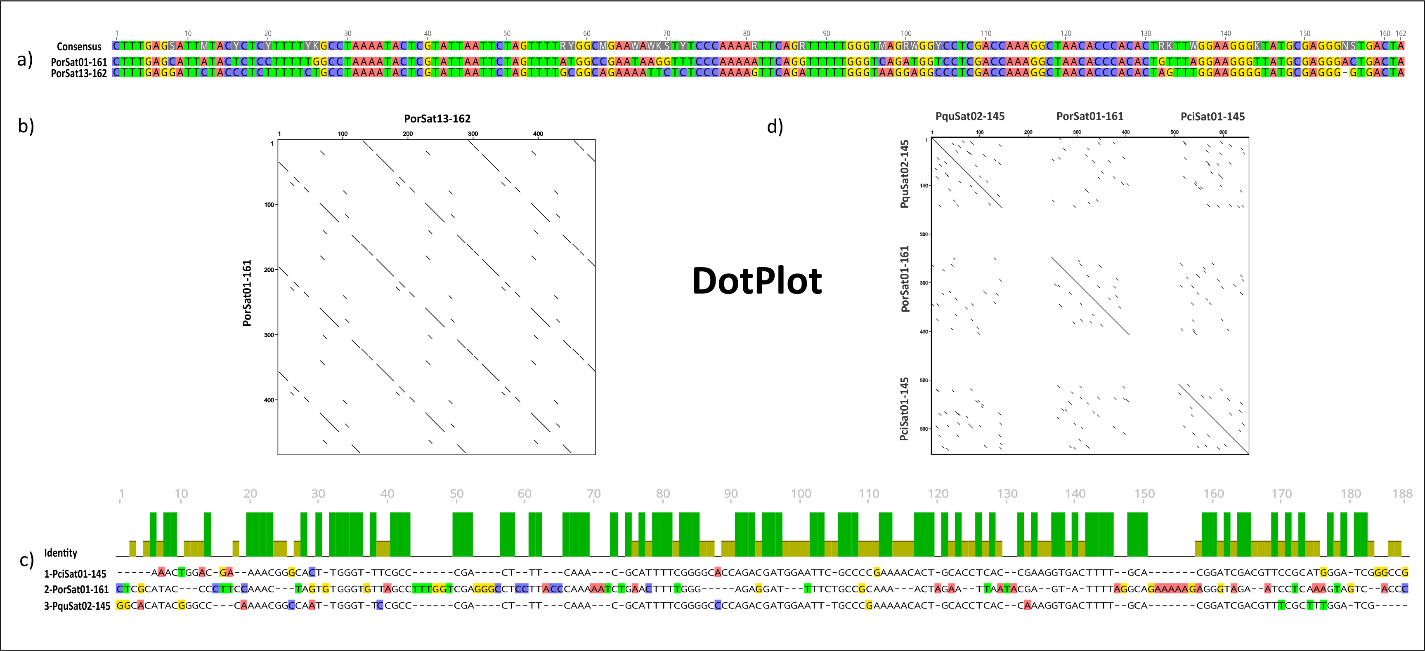
**Online Resource 1.** Similarity analysis of *Passiflora* superfamily 1 satDNA. a) Alignment and b) dotplot of satDNAs PorSat01-161 and PorSat13-162 sequences, showing a high similarity (84% identity) between them. c) Alignment and d) dotplot of satDNA PorSat01-161, PquSat02-145 and PciSat01-145 sequences showing 54% identity.

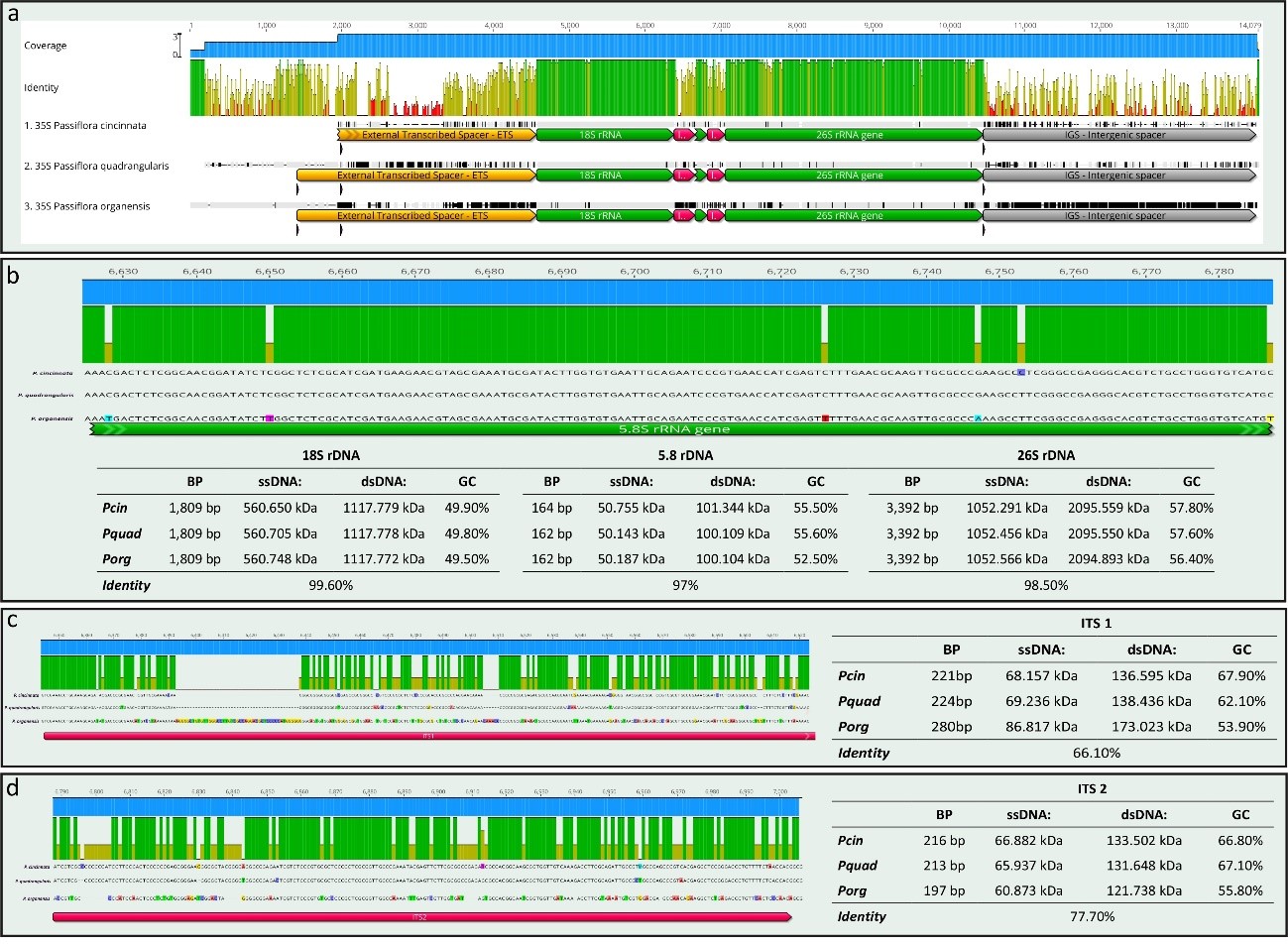
**Online Resource 2.** Alignment of 35S ribosomal DNA between *P. organensis*, *P. cincinnata* and *P. quadrangularis.* Details from b) rDNA genes c) ITS 1 and d) ITS 2, showing different comparative parameters: size (bp), molecular weight (ssDNA) and dsDNA, and GC content.

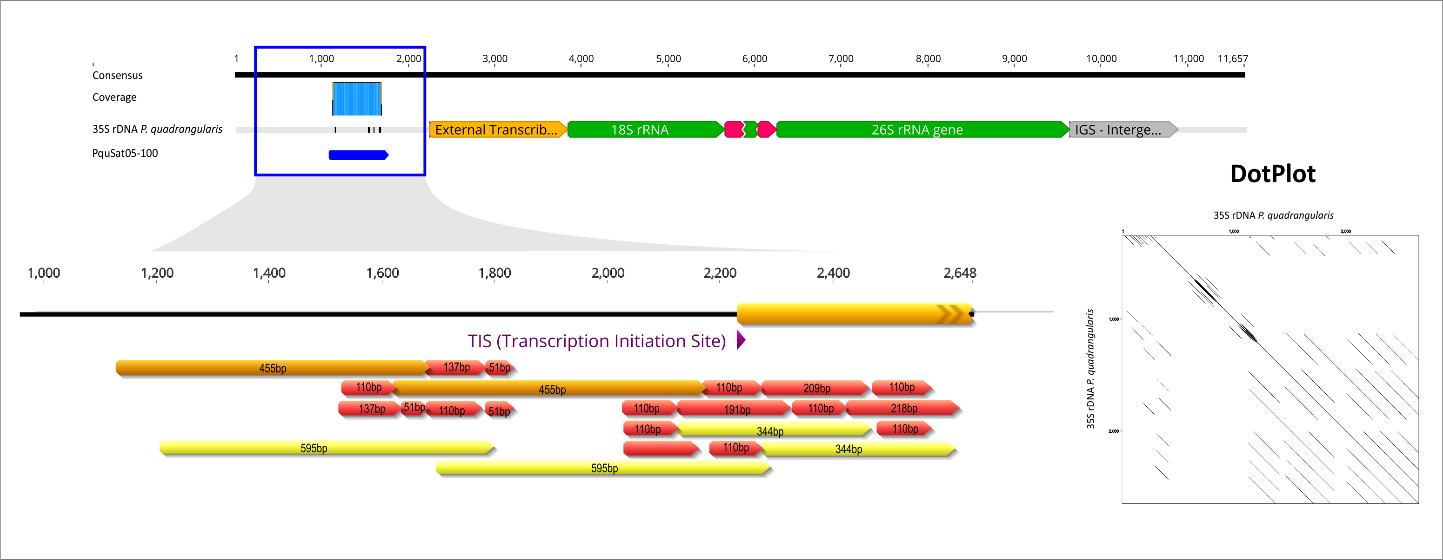
**Online Resource 3.** Organization of the 35S ribosomal DNA (rDNA) sequence of *Passiflora quadrangularis* assembled by NOVOPlasty. Comparison of the sequence similarity between the consensus sequence of the satDNA PquSat01-100 and the intergenic spacer (IGS) region of 35S rDNA, showing detail of repeats and subrepeats and the dotplot analysis of the complex structure of satDNA PquSat01-100.
